# Metagenome diversity illuminates origins of pathogen effectors

**DOI:** 10.1101/2023.02.26.530123

**Authors:** Victoria I. Verhoeve, Stephanie S. Lehman, Timothy P. Driscoll, John F. Beckmann, Joseph J. Gillespie

## Abstract

Recent metagenome assembled genome (MAG) analyses have profoundly impacted Rickettsiology systematics. Discovery of basal lineages (Mitibacteraceae and Athabascaceae) with predicted extracellular lifestyles reveals an evolutionary timepoint for the transition to host dependency, which occurred independent of mitochondrial evolution. Notably, these basal rickettsiae carry the Rickettsiales *vir* homolog (*rvh*) type IV secretion system (T4SS) and purportedly use *rvh* to kill congener microbes rather than parasitize host cells as described for derived rickettsial pathogens. MAG analysis also substantially increased diversity for genus *Rickettsia* and delineated a basal lineage (*Tisiphia*) that stands to inform on the rise of human pathogens from protist and invertebrate endosymbionts. Herein, we probed Rickettsiales MAG and genomic diversity for the distribution of *Rickettsia rvh* effectors to ascertain their origins. A sparse distribution of most *Rickettsia rvh* effectors outside of Rickettsiaceae lineages indicates unique *rvh* evolution from basal extracellular species and other rickettsial families. Remarkably, nearly every effector was found in multiple divergent forms with variable architectures, illuminating profound roles for gene duplication and recombination in shaping effector repertoires in *Rickettsia* pathogens. Lateral gene transfer plays a prominent role shaping the *rvh* effector landscape, as evinced by the discover of many effectors on plasmids and conjugative transposons, as well as pervasive effector gene exchange between *Rickettsia* and *Legionella* species. Our study exemplifies how MAGs can provide incredible insight on the origins of pathogen effectors and how their architectural modifications become tailored to eukaryotic host cell biology.

## INTRODUCTION

Until recently, Order Rickettsiales (*Alphaproteobacteria*) contained three families harboring diverse obligate intracellular parasites ^1^. Rickettsiaceae and Anaplasmataceae are best studied and harbor invertebrate endosymbionts, human pathogens, and reproductive parasites ^2–7^. Midichloriaceae contains some arthropod-associated bacteria of unknown vertebrate pathogenicity ^8^, but most species are described from protists ^9–14^. Remarkably, Castelli and colleagues ^15^ described the first extracellular rickettsial species, “*Candidatus* Deianiraea vastatrix”, as a bacterium dependent on *Paramecia* and sharing many characteristics of the intracellular lifestyle. A new family, Deianiraeaceae, was proposed, calling into question the specific timepoint in rickettsial evolution wherein obligate intracellularity emerged from an obligate extracellular or facultative intracellular lifestyle.

Historically, Rickettsiales were widely considered a sister lineage to the mitochondria progenitor, with this assemblage representing a basal branch of the *Alphaproteobacteria* ^16–20^. Pioneering work on rickettsial genomes supported this assertion, identifying decreased genome size and pseudogenization of genes within many metabolic pathways, conceptualized as “reductive genome evolution”, behind addiction to the eukaryotic cytosol ^18, 21–25^. This dogma held for two decades as hundreds of diverse Rickettsiales genomes were sequenced ^6, 26^. However, phylogenetic analysis of deep marine **metagenome-assembled genomes** (**MAGs**) illustrated that mitochondria likely originated outside of all described *Alphaproteobacteria* ^27^, and recent phylogenomic description of certain novel MAGs established two basal rickettsial lineages, Mitibacteraceae and Athabascaceae, with features indicating an extracellular lifestyle not dependent on eukaryotic hosts ^28^. These findings bolstered the growing trend for identifying mostly aquatic, protist-associated rickettsial species with traits (e.g., flagella, larger genome size, greater metabolic capacity, etc.) more characteristic of free-living and facultative intracellular bacteria but absent from the numerous genomes of well-characterized invertebrate- and animal associated rickettsial species ^13, 15, 29–33^. Furthermore, they establish a Rickettsiales phylogenetic framework that allows for assessing the evolutionary trajectories within five derived lineages for innovations that emerged from transitions to host dependency.

Estimated to have arisen ∼1.9 billion years ago ^34^, the *Alphaproteobacteria* is highly diversified in form and function yet rife with convergence in morphology and lifestyle through common adaptation to numerous environments, including eukaryotic cells ^35^. Order-level signatures are rare, making it curious that the genomes of Mitibacteraceae and Athabascaceae encode the **Rickettsiales *vir* homolog** (***rvh***) **type IV secretion system** (**T4SS**), a Rickettsiales signature that functions in overtaking eukaryotic cells ^36–40^. The *rvh* T4SS is odd in its design ^41, 42^, with specialized duplications of some components hypothesized to autoregulate effector secretion ^43, 44^. Effectors have been characterized for species of *Ehrlichia* ^45–48^, *Anaplasma* ^49–54^, and *Rickettsia* ^55–57^. Schön et al. proposed that species of Mitibacteraceae and Athabascaceae utilize the *rvh* T4SS for killing congener microbes ^28^, provided that these genomes harbor candidate *rvh* effectors with characteristics similar to effectors in other T4SS and T6SS killing machines ^58, 59^. Thus, the five derived families purportedly repurposed the *rvh* T4SS to secrete effectors that commandeer host cellular processes to support intracellular replication (or epicellular parasitism in the case of Deianiraeaceae species).

The existence of an ancient secretion machine, yet independent gain of its effectors later in evolution, prompted us to poll the ever-growing MAG diversity for clues on *rvh* effector origins. We focus on known or candidate effectors from the genus *Rickettsia*, as recent studies have considerably expanded Rickettsiaceae diversity. Genome sequences from “environmental” Rickettsiaceae species (i.e. those from protists, apicomplexans, diplomonads, crustaceans, and insects) have illuminated basal lineages of Rickettsiaceae that are critical for inferring the emergence of genomic traits in *Orientia* and *Rickettsia* pathogens ^29, 60–64^. Further, phylogenetic analysis of genome sequences from novel genera “*Candidatus* Sarmatiella” (paramecium symbiont) ^65^ and “*Candidatus* Megaira” (symbionts of algae and ciliates) ^31, 66^ indicate *Orientia* and *Rickettsia* species are more divergent than previously appreciated. Finally, a long-standing recognized basal lineage of *Rickettsia* termed “Torix Group”, which is highly diverse and widely present in non-blood-feeding arthropods ^67–71^, was recently classified as a new genus, “*Candidatus* Tisiphia”, in a study that identified many new provisional *Rickettsia* (and *Tisiphia*) species from MAG analyses of diverse arthropods ^72^.

We present phylogenomic analyses that effectively demonstrate the utility of MAG data for not only inferring the origins of pathogen effectors, but for better understanding effector architectures through enhanced predictive power from greater sequence diversity. Provided that many MAGs come from environmental sampling or eukaryotes with no known human association, our approach stands to inform on the evolution of vertebrate pathogenesis.

## RESULTS AND DISCUSSION

### Mapping the Acquisition of Rickettsial Effectors

We hypothesize that the transition to an intracellular lifestyle necessitated the acquisition of a more diverse effector repertoire. Thus, to gain an appreciation of the origins and conservation of *Rickettsia rvh* T4SS effectors, we performed a phylogenomics analysis encompassing the newly appreciated rickettsial diversity (**FIG. 1A**). This initially involved creating a matrix of taxa (genomes and metagenomes) determined to encode the *rvh* T4SS and the distribution of effectors. Six ***rvh* effector molecules** (**REMs**: RalF, Pat1, Pat2, Risk1, RT0527, and RARP-2) and 14 **candidate REMs** (**cREMs**) were evaluated based on prior studies implicating their secretion and/or interaction with the *rvh* coupling protein (RvhD4), or presence of motifs known to target either congener bacteria or eukaryotic molecules ^55–57, 73–80^ (**FIG. 1B**). Our analyses expanded two REMs (Pat1 and RARP-2) and four cREMs based on identification of duplications (cREM-1, cREM-2, cREM-4), a partner protein (cREM-5), and a domain within the **surface cell antigen** (**sca**) Sca4 that we demonstrate to be widespread in non-Sca4 proteins. Collectively, a total of 26 proteins were analyzed within the phylogenomic framework (**FIG. 1C**; **FIG. 2**). A phylogeny was estimated from concatenated alignments of RvhB4-I and RvhB4-II proteins from 153 genome assemblies (**FIG. 1C**; **FIG. 2**; **Table S1**; **FIG. S2**). This collective matrix is an effective framework for mapping the earliest occurrence of these *rvh* effectors in the rickettsial tree, additionally identifying several likely origins for **lateral gene transfer** (**LGT**).

**FIGURE 1.**
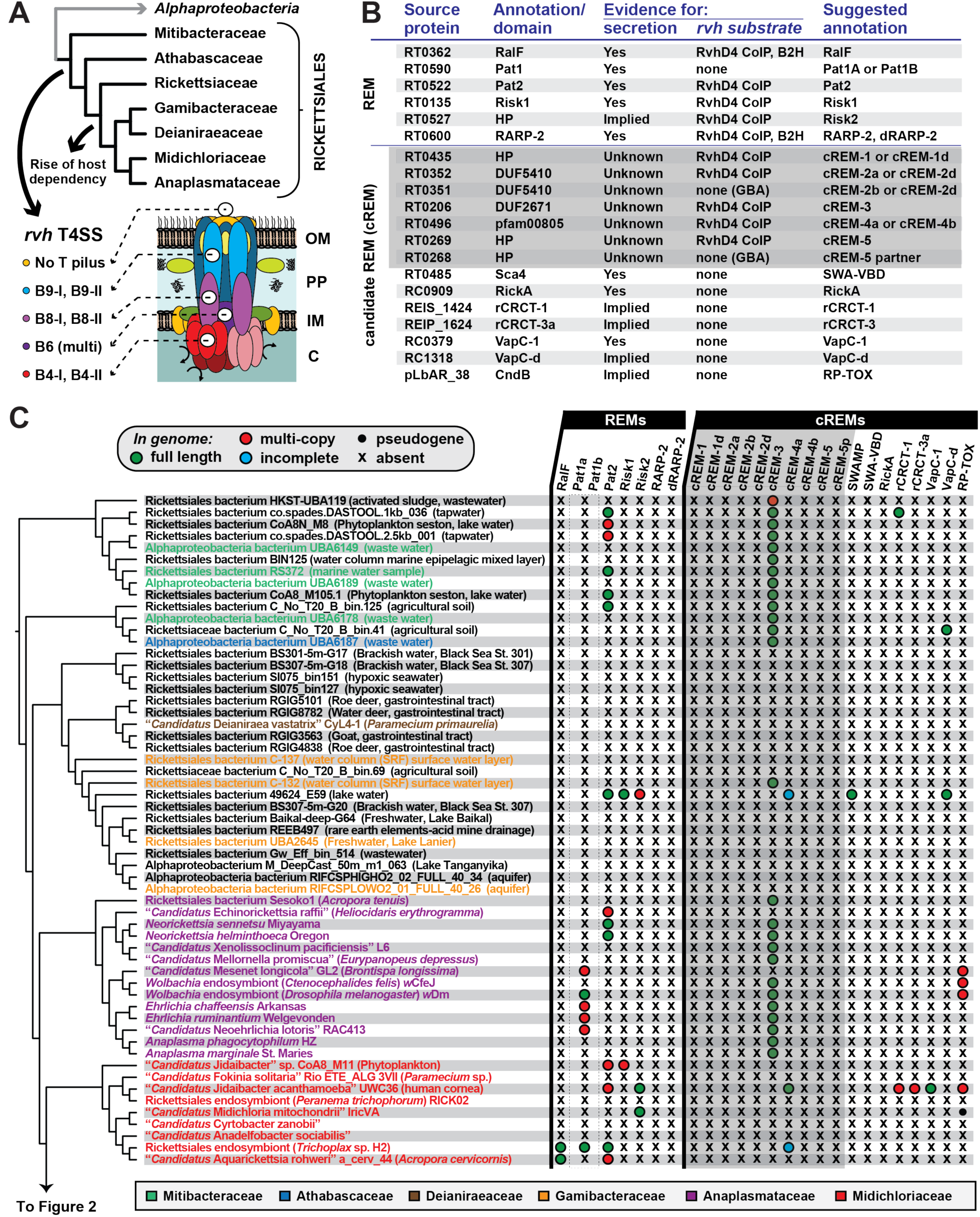
Probing Rickettsiales diversity for the evolution of *Rickettsia* type IV secretion system effectors. **(A)** The atypical Rickettsiales *vir* homolog (*rvh*) T4SS is a hallmark of Rickettsiales, arising before the origin of host dependency ^28, 197^. Schema depicts recent genome-based phylogeny estimation ^28^. Rvh characteristics ^37, 44^ are described at bottom and further in FIG. S1. **(B)** List of *Rickettsia rvh* effector molecules (REMs) and candidate REMs (cREMs). GBA, guilty by association. ‘Implied’ means analogous proteins are known to be secreted by other bacteria and/or the effector has strongly predicted eukaryotic molecular targets. Secretion, coimmunoprecipitation (CoIP), and bacterial 2-hybrid (B2H) data are compiled from prior reports ^55–57, 73–80^. SWA, Schuenke Walker Antigen domain. **(C)** Phylogenomic analysis of *Rickettsia* REMs and cREMs in non-Rickettsiaceae lineages. Cladogram summarizes a phylogeny estimated from concatenated alignments for RvhB4-I and RvhB4-II proteins from 153 rickettsial assemblies (full tree, **FIG. S2**; sequence information, **Table S1**). Non-Rickettsiaceae lineages are shown (see **FIG. 2** for Rickettsiaceae). Dashed box for Pat1 proteins indicates the inability to discern Pat1a and Pat1b homology outside of *Tisiphia* and *Rickettsia* species (see **FIG. 2**; **FIG. 4**). SWAMP, SWA modular proteins. Information for all REMs and cREMs is provided in **Table S2**.

**FIGURE 2.**
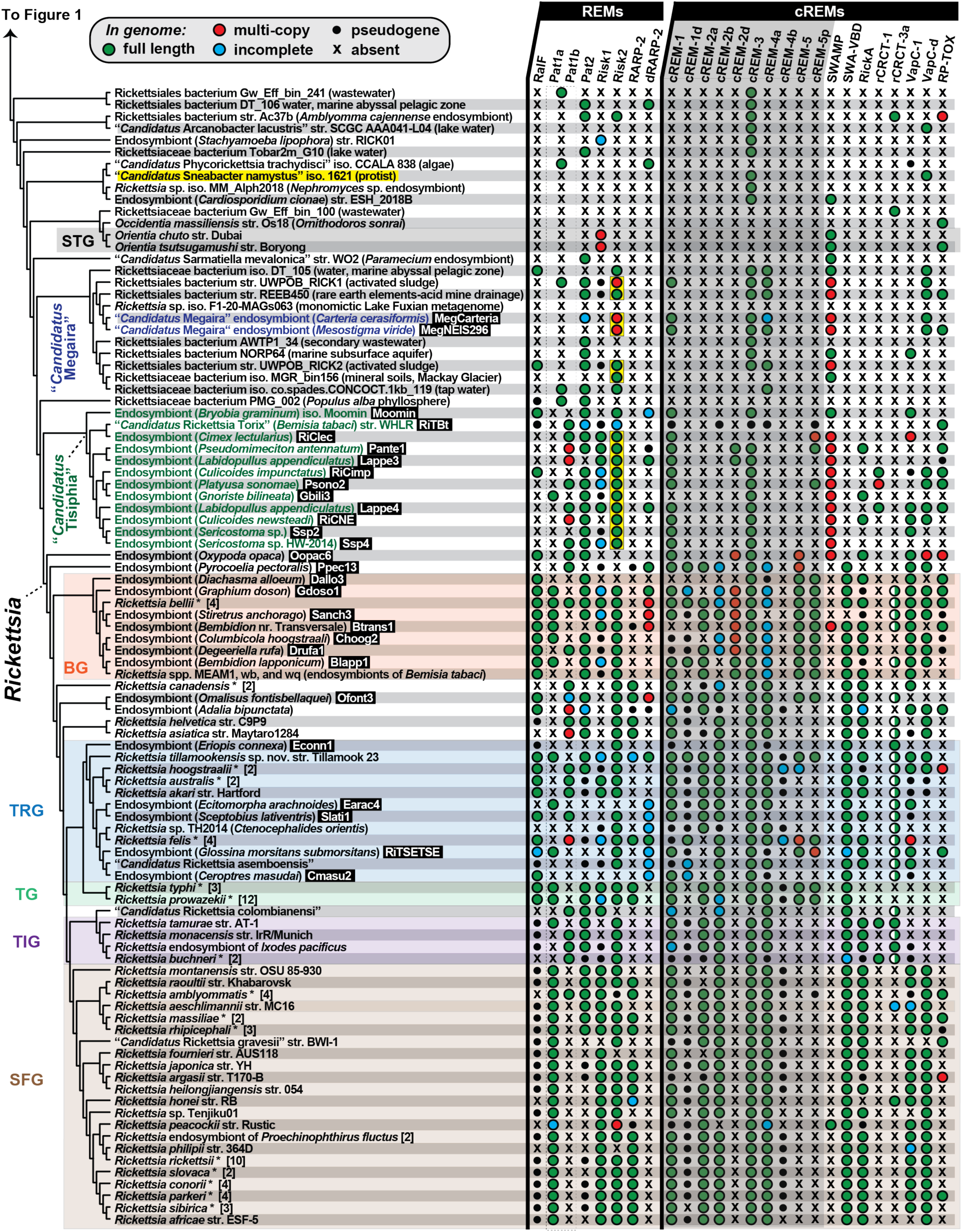
Phylogenomic analysis of *Rickettsia* REMs and cREMs in Rickettsiaceae. Cladogram (continued from FIG. 1C) summarizes a phylogeny estimated from concatenated alignments for RvhB4-I and RvhB4-II proteins from 153 rickettsial assemblies (full tree, FIG. S2; sequence information, **Table S1**). “*Candidatus* Sneabacter namystus” (highlighted yellow) was manually added to the cladogram based on prior phylogeny estimation ^184, 185^ as this species lacks *rvh* genes but carries a type VI secretion system T6SS (see **FIG. S3**). Black boxes provide short names for 29 MAGs from Davison *et al*. ^72^ (NOTE: the clade colored green comprises genus *Tisiphia* though genus name *Rickettsia* reflects NCBI taxonomy as of Feb. 26^th^, 2023). Asterisks depict multiple genome assemblies for a species. BG, Bellii Group; TRG, Transitional Group; TG, Typhus Group; TIG, Tamurae-Ixodes Group; SFG, Spotted Fever Group. Dashed box for Pat1 proteins indicates the inability to discern Pat1a and Pat1b homology outside of *Tisiphia* and *Rickettsia* species (see **FIG. 4**). Yellow boxes denote Risk2 proteins that are appended to C-terminal Schuenke-Walker antigen (SWA) domains (see **FIG. 5**). SWAMP, SWA modular proteins; all other REMs and cREMs are described in **FIG. 1B** and **Table S2**). Half circles for rCRCT-3a depict the presence of one or more antidotes but no toxin.

### Origins of REMs

#### An emerging diversity of bacterial Arf-GEFs

Bacterial mimicry of eukaryotic-like **Sec7 domains** (**S7D**) to function as **guanine nucleotide exchange factors** (**GEF**) for host **ADP-ribosylation factors** (**Arfs**) was first described for *Legionella pneumophila*, which utilizes the *dot*/*icm* T4SS effector RalF to recruit and activate host Arf1 to the ***Legionella*-containing vacuole** (**LCV**) ^81, 82^. Certain *Rickettsia* genomes encode RalF proteins that are remarkably similar to *Legionella* counterparts across the S7D, as well as a **Sec7 capping domain** (**SCD**) that restricts access to the catalytic site ^83–85^. The SCD has high specificity for host membranes and differentially regulates effector subcellular localization for *Legionella* (the LCV) and *Rickettsia* (the inside face of plasma membrane) RalF ^86^. *Rickettsia* RalF was the first characterized REM; its secretion during host cell invasion activates host Arf6 at the plasma membrane, a process driven by a unique C-terminal extension, termed **Variable with Pro-rich region** (**VPR**), that interacts with host actin and **phosphatidylinositol 4,5-biphosphate** (**PI(4,5)P_2_**) at entry foci ^55, 87^. The presence of *ralF* in the genomes of some *Rickettsia* pathogens but absence in non-pathogenic species led to speculation that this REM may be a lineage specific virulence factor ^55, 87, 88^. Further, while species of *Rickettsia* and *Legionella* exchange genes in common intracellular environments ^89, 90^, the absence of *ralF* in any other known bacteria precluded insight on the origin of RalF and specifically how the diverse C-terminal architectures arose.

Our analyses provide clarity on RalF evolution by unearthing numerous bacterial analogs with novel S7D-containing architectures (**FIG. 3**; **FIG. S4**). First, an unusual *Legionella* RalF from *L*. *clemsonensis* was found to carry a conserved domain at its C-terminus that was also detected in a large **ankyrin** (**ANK)** repeat-containing protein of the *Rickettsia* endosymbiont of *Graphium doson* (Gdoso1) genome (**FIG. 3A**; **FIG. S4A,B**). This Gdoso1 protein also contains another conserved domain at its C-terminus that is widespread in *Rickettsia* genomes. These observations indicate frequent recombination in conjunction with LGT of these diverse genes. Second, while the most basal *Rickettsia* species (endosymbiont of *Oxypoda opaca*, “Oopac6”) harbors a RalF with the C-terminal VPR, two *Tisiphia* species (endosymbionts of *Bryobia graminum*, or “Moomin”, and *Culicoides impunctatus*, or “RiCimp”) are *Legionella*-like in C-terminal domain architecture (**FIG. 3A**,**C**; **FIG. S4C-E**). RiCimp RalF is encoded on a plasmid (pRiCimp001), the first for a *ralF* gene (**FIG. 3D**), supporting original speculation for RalF exchange between *Legionella* and *Rickettsia* species ^81^. Remarkably, pRiCimp001 also carries a **toxin-antidote** (**TA**) module highly similar to the plasmid-encoded TA module of *Rickettsia felis* str. LSU-Lb ^91^ that we previously described as part of the mobilome shuttling **reproductive parasitism** (**RP**) genes across *Rickettsia* and wolbachiae ^91^ (discussed below, section

**FIGURE 3.**
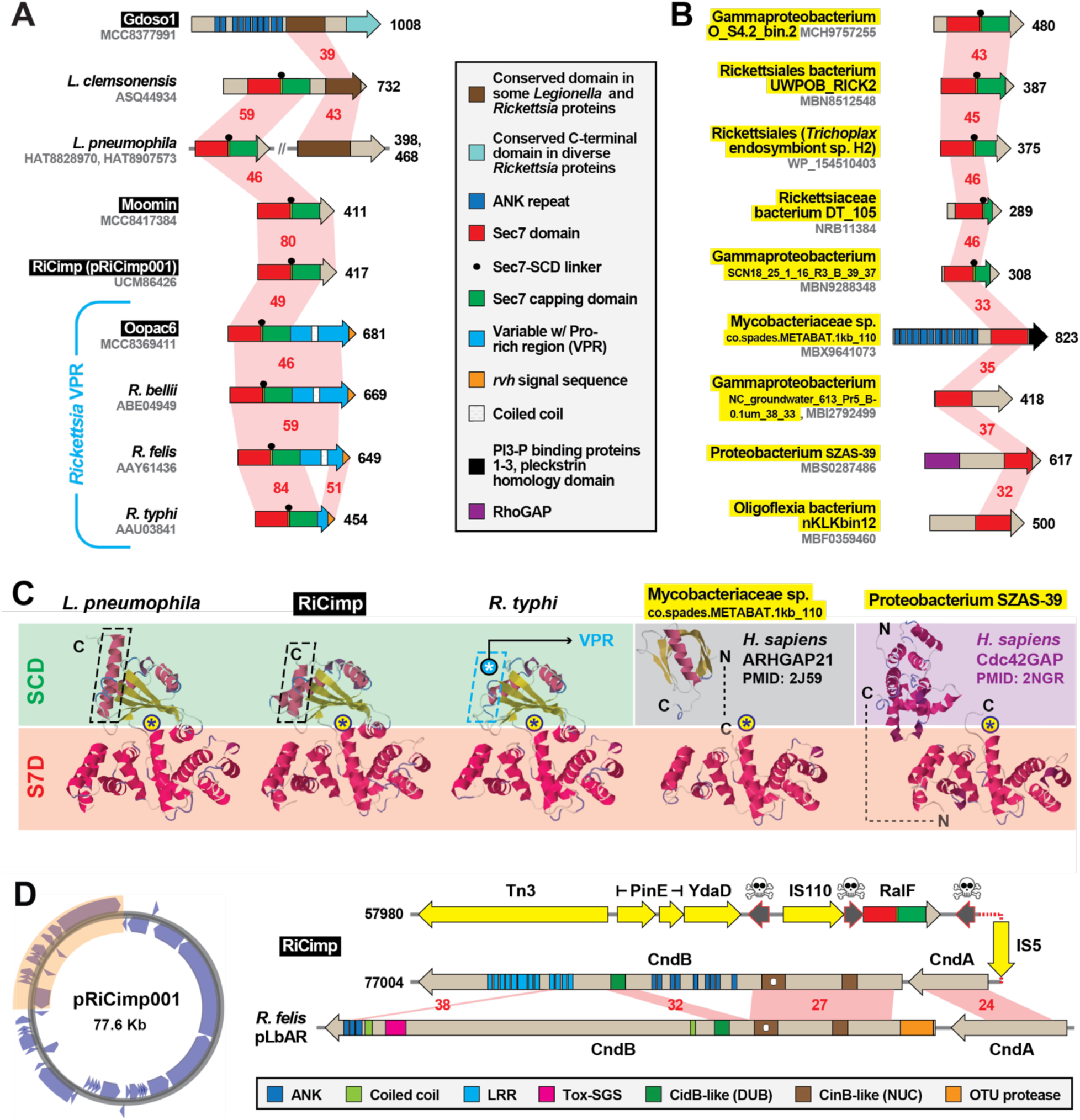
MAG analysis divulges a greater diversity of Sec7-domain-containing proteins than previously appreciated. Black boxes provide short names for MAGs from Davison *et al*. ^72^. These and additional newly discovered RalF-like proteins (highlighted yellow) substantially expand the prior RalF diversity. Structural models for proteins are found in FIG. S4D-F. **(A**,**B)** Novel insight from new **(A)** *Legionella* and rickettsial architectures and **(B)** new RalF-like proteins discovered in MAGs. Red shading and numbers indicate % aa identity across pairwise alignments (sequence information in **Table S2**). All protein domains are described in the gray inset. **(C)** Comparison of the *Legionella pneumophila* RalF structure (PDB 4C7P) ^85^ with predicted structures of S7D-SCD regions of RiCimp RalF (LF885_07310) and *R*. *typhi* RalF (RT0362), and S7Ds of Mycobacteriaceae sp. co.spades.METABAT.1kb_110 (K2X97_15435) and Proteobacterium SZAS-39 (JSR17_09325). The delineation of the Sec7 domain (S7D, red) and Sec7-capping domain (SCD, green if present), is shown with an approximation of the active site Glu (asterisk). Additional eukaryotic-like domains for the non-rickettsial proteins are noted. Modeling done with Phyre2 ^192^. More detailed structural explanation is found in FIG. S4C. **(E)** RiCimp plasmid pRiCimp001 carries RalF and a TA module similar to those characterized in reproductive parasitism^91^. Gene region drawn to scale using PATRIC compare region viewer tool ^198^. Yellow, transposases and other mobile elements; skull-and-crossbones, pseudogenes; other domains described in gray inset at bottom. Plasmid map created with Proksee (https://proksee.ca/).

**FIGURE 4.**
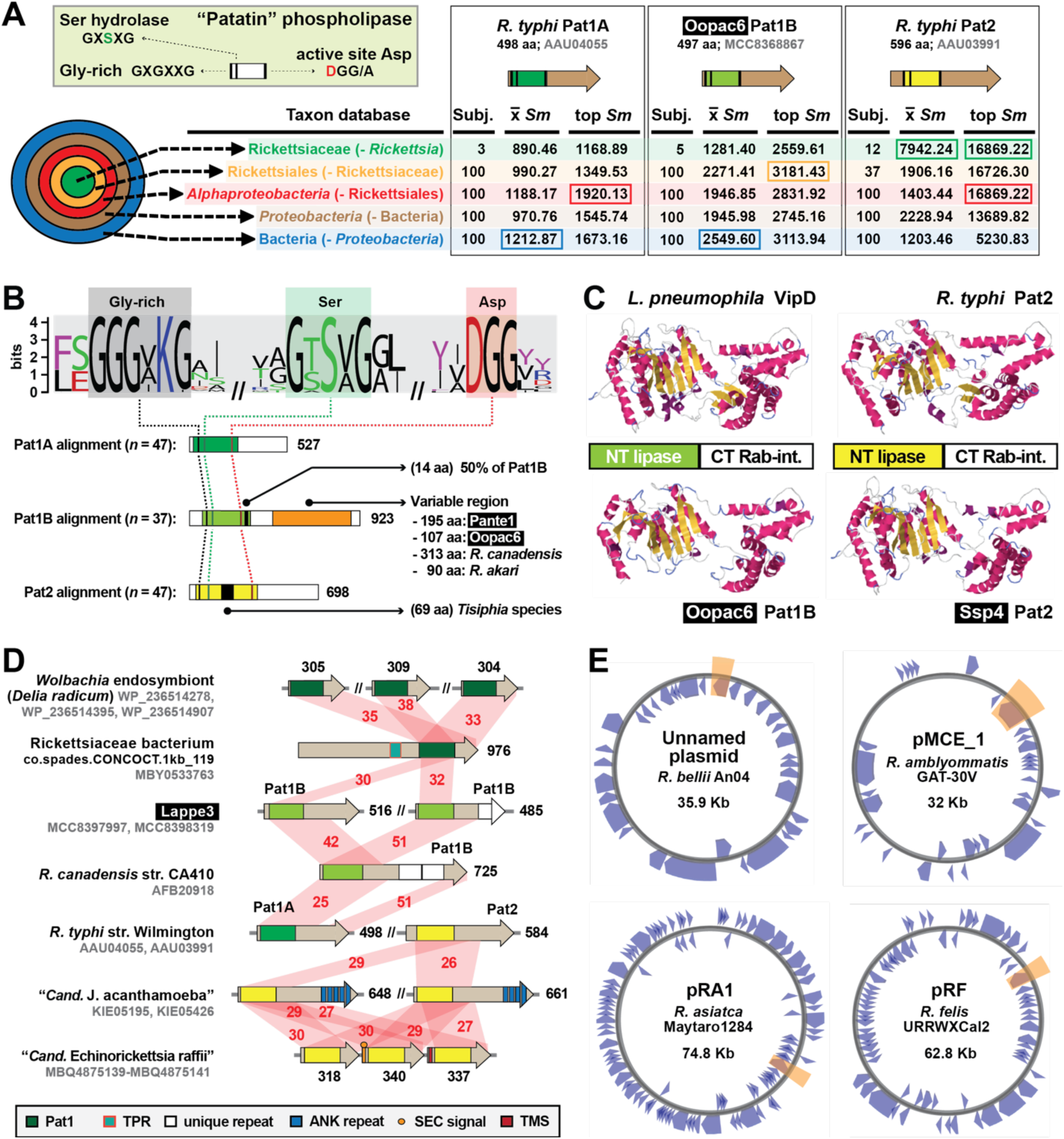
Divergent patatin phospholipases are recurrent in rickettsial evolution. Black boxes provide short names for MAGs from Davison *et al*. ^72^. **(A)** PLA_2_ active site characteristics and divergent patatin forms. Green inset describes general patatin domain and active site architecture. HaloBlast results for Pat1A, Pat1B, and Pat2 (query sequences described at top) are shown, with top-scoring halos boxed (full results in **TABLE S4**). **(B)** Sequence logo ^187^ showing conservation of the PLA_2_ active site motifs across *Tisiphia* and *Rickettsia* patatins (sequence information provided in **Table S2**). Pat1A, Pat1B, and Pat2 sequences were aligned separately with MUSCLE ^186^ (default parameters) with active site motifs compiled for conservation assessment. Features unique to each patatin are noted. **(C)** *Legionella pneumophila* VipD structure (PDBID: 4AKF) and modeling of three rickettsial patatins to VipD using Phyre2 ^192^. **(D)** Diverse architectures for select patatins. Red shading and numbers indicate % aa identity across pairwise alignments (sequence information in **Table S2**). All protein domains are described in the gray inset. Dark green indicates Pat1 domains not grouped into A or B. **(E)** Four *Rickettsia* plasmids carry *pat1B* (shaded orange). Plasmid maps created with Proksee (https://proksee.ca/).

#### cREMs with Characterized Function)

Finally, RalF proteins from three additional rickettsial species and two putative gammaproteobacterial species carry both the S7D and SCD but no C-terminal extensions (**FIG. 3B**; **FIG. S4E,F**). Four other S7D-containing proteins from non-rickettsial bacteria lack SCDs; however, two contain eukaryotic domains found in **RHO GTPase-activating proteins** (**RHOGAP**) that also target Arfs (**FIG. 3B**,**C**; **FIG. S4F**). All of these collective proteins have a highly conserved S7D and SCD (if present) and include most of the structural features that define RalF proteins (**FIG. S4G**). These collective characteristics attest to LGT disseminating the S7D-SCD architecture across divergent bacteria, with recurrent gains of additional domains tailored to eukaryotic cell functions (e.g., VPR, ANK, and RHOGAP). We posit that the acquisition of the *Rickettsia*-unique C-terminal VPR occurred early in *Rickettsia* evolution, with multiple losses of RalF in more than half of the sequenced species (**FIG. 2**).

#### Patatins: divergent phospholipases are recurrent in Rickettsiales

Rickettsiae (and other Rickettsiales species lysing the phagosome and/or lysing host cells) require membranolytic effectors throughout the intracellular lifestyle. **Phospholipase D** (**PLD**) is a highly conserved enzyme with demonstrated membranolytic activity in a surrogate expression system ^92^ though an unresolved function during *Rickettsia* infection of host cells ^93^. PLD contains a N-terminal Sec signal ^88^, yet other **phospholipase A_2_** (**PLA_2_**) enzymes (patatins Pat1 and Pat2) have sequence characteristics of *rvh* substrates ^74^ and Pat2 binds RvhD4 in coimmunoprecipitation assays ^56^ (**FIG. 1B**). Studies on *R*. *typhi* have shown that both Pat1 and Pat2 are secreted during host cell infection, require host cofactors for activation, and function early in infection by facilitating phagosome escape ^74, 75^. Recent work on *R*. *parkeri* Pat1 also demonstrated a role in phagosome escape in addition to facilitating avoidance of host polyubiquitination and autophagosome maturation, as well as promotion of actin-based motility and intercellular spread ^94^. *R*. *parkeri* lacks Pat2, which is slightly more restricted in *Rickettsia* genomes and possibly provides a function in host cell lysis for rickettsiae that do not spread intercellularly without host cell lysis (e.g., TG rickettsiae).

All patatins share a common active site architecture that is critical for PLA_2_ activity (**FIG. 4A**). Despite this, Pat1 and Pat2 are highly divergent outside of the patatin domain and have different origins based on phylogeny estimation ^74^. Further, Pat1 proteins form two distinct groups, Pat1A and Pat1B, with *pat1B* found on plasmids and often recombining with chromosomal *pat1* loci ^74^. Utilizing newly discovered rickettsial patatins from MAGs, we show that all three enzymes (Pat1A, Pat1B, and Pat2) have distinct sequence profiles, with Pat1B proteins having a highly length variable C-terminal region relative to Pat1A and Pat2 enzymes (**FIG. 4A**,**B**). Despite this, Pat1B and Pat2 proteins have cryptic similarity across their C-terminal regions to support robust modeling to the crystal structure of *L*. *pneumophila dot*/*icm* T4SS effector VipD ^95^ (**FIG. 4C**). During *L*. *pneumophila* host cell infection, secreted VipD localizes to host endosomes, catalyzing the removal of **phosphatidylinositol 3-phosphate** (**PI(3)P**) from endosomal membranes (N-terminal patatin domain) and binding Rab5 or Rab25 (C-terminal domain), ultimately blocking endosome-LCV fusion ^95–97^. As with RalF, it is likely that *Rickettsia* Pat1 and Pat2 proteins have rudimentary analogous functions to VipD (targeting host membranes and binding host Rabs) but spatiotemporal and biochemical differences provided that rickettsiae lyse the phagosome and seemingly do not interfere with early endosome trafficking on par with *Legionella* species.

We detected Pat1 and Pat2 proteins in several non-Rickettsiaceae genomes (**FIG. 1C**), with some genomes (e.g., novel sea urchin and cabbage root fly endosymbionts in the Anaplasmataceae ^98, 99^) harboring duplications (**FIG. 4D**). Pat1 proteins from Rickettsiales species outside of the genera *Tisiphia* and *Rickettsia* could not be confidently assigned to either Pat1A or Pat1B (**FIG. 1C**; **FIG. 2**; dark green domains in **FIG. 4D**). Most species of *Tisiphia* and *Rickettsia* carry either Pat1A or Pat1B and/or Pat2 (**FIG. 2**). The only two species carrying all three distinct enzymes (*R*. *bellii* and *R*. *amblyommatis*) have Pat1B encoded on a plasmid (**FIG. 4E**). Overall, the patchwork distribution of these divergent enzymes, evidence for modular domain diversification, and presence on plasmids indicate that PLA_2_ activities for rickettsiae are lineage-specific and subject to continual patatin gene gain and loss throughout evolution. Further, certain *pat* gene profiles may confer advantages in particular hosts.

#### Domain repurposing is Risky business

Bacterial pathogens can directly modify host membrane **phosphaditylinositol** (**PI**) composition by mimicking eukaryotic kinases, phosphatases, and phosphotransferases ^100–103^. Secreted PI kinases from intracellular pathogens *R*. *typhi* (Risk1), *L*. *pneumophila* (LegA and LepB), and *Francisella tularensis* (OpiA) alter the PI composition on phagosomes to prohibit maturation and fusion with lysosomes ^56,104,105^. Characterized as either PI3 (Risk1, LegA, and OpiA) or PI4 (LepB) kinases, these enzymes possess a similar PI3/4 active site architecture (pfam00454) analogous to eukaryotic PI kinases, as well as certain protein kinases, that function in a myriad of membrane-associated functions, including intracellular signalling and trafficking ^106^ (**FIG. 5A**,**B**). Subverting these host cell functions is highly advantageous to intracellular parasitism; thus, the dearth of identified PI3/4 kinase effectors likely reflects the cryptic nature of the PI3/4 active site within these proteins, which lack similarity outside of the PI3/4 active site domain ^56, 105^.

**FIGURE 5.**
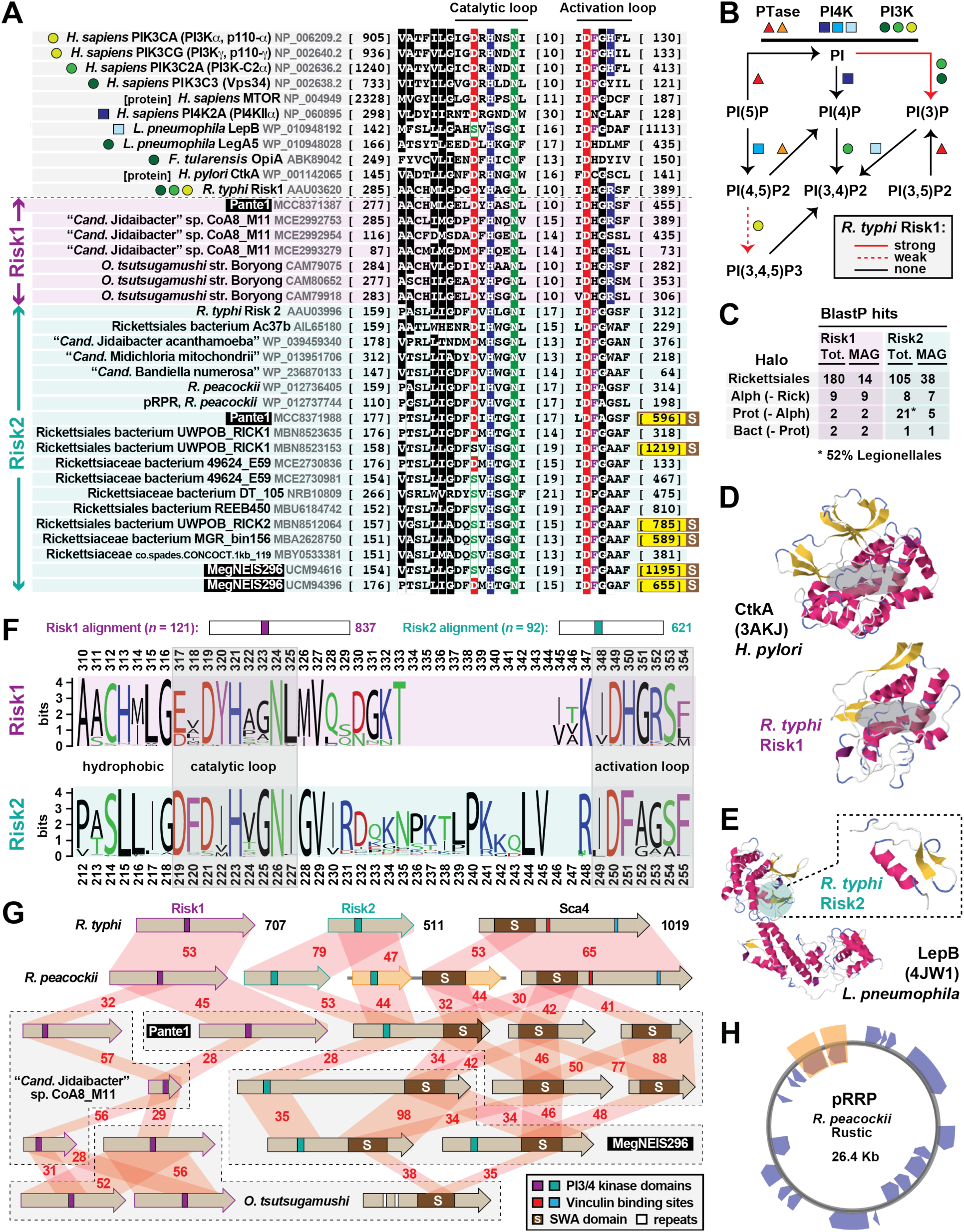
Discovery of a novel rickettsial PI kinase that can associate with a widespread rickettsial surface antigen. Black boxes provide short names for MAGs from Davison *et al*. ^72^. Amino acid coloring is described in the **FIG. 3** legend. **(A)** Previous work (above dashed line) identified a *Rickettsia* PI kinase, Risk1, with a cryptic active site similar to human and other bacterial PI3/PI4 kinases and related protein kinases. Colored shapes depict characterized substrate specificity (see panel **B**). Our study (below dashed line; select proteins shown) identified new rickettsial Risk1 proteins, as well as a second PI kinase (Risk2) also prevalent in rickettsial genomes and MAGs (FIG. 1C; **FIG. 2**). All PI3/PI4 kinase domains were aligned using MUSCLE ^186^ (default parameters). Sequence information is provided in **Table S2**. Yellow highlighting on end coordinates denotes Risk2 proteins fused to a C-terminal SWA domain (see panel **G**; FIG. S5). **(B)** Mechanisms of phosphorylation on the PI inositol ring at 3’, 4’ and 5’ positions. Data for *R*. *typhi* Risk1 is superimposed in red ^56^. **(C)** HaloBlast results (*R*. *typhi* Risk1 and Risk2 as queries) broken down to illustrate the presence of *Rickettsia*-like PI kinases in MAGs and the similarity between Risk2 and *Legionella* PI kinases (full data in **Table S2**). **(D)** Risk1 threads with high confidence (90.7%, 72% coverage; Phyre2 ^192^) to the *Helicobacter pylori* proinflammatory kinase CtkA ^108^. **(E)** Risk2 threads with high confidence (85.1%, 9% coverage; Phyre2 ^192^) to a limited region of LepB, a Rab GTPase-activating protein effector from *L. pneumophila* ^199^. **(F)** Risk1 and Risk2 proteins have cryptic and distinct PI3/4 active sites, yet lack similarity outside of these regions. Logos depict individual alignments, which are summarized at top and were performed with MUSCLE ^186^, default settings. **(G)** Rickettsiae utilize a diverse arsenal of PIK effectors, some of which are tethered to SWA domains. Six select species are shown with their full complement of PI3/4 kinase and SWA architectures. Red shading and numbers indicate % aa identity across pairwise alignments (sequence information in **Table S2**). *R. peacockii* pRRP proteins are shaded orange (see panel **H**). **(H)** Plasmid pRRP of *R. peacockii* str. Rustic ^112, 200^ carries a divergent Risk2 gene that is adjacent to an ORF encoding a SWAMP (orange highlighting). Plasmid map created with Proksee (https://proksee.ca/).

BlastP and HMMER ^107^ analyses using only the Risk1 PI3/4 active site revealed nearly 300 Rickettsiales proteins, with many genomes having multiple divergent kinases. Further inspection revealed the presence of a second conserved protein harboring the PI3/4 active site, which we named ***Rickettsia* intracellular secreted kinase-2** (**Risk2**) (**FIG. 5A**). Notably, the *R*. *typhi* protein (RT0527) was captured in the same RvhD4 coimmunoprecipitation assay that identified Risk1 as a REM ^56^ (**FIG. 1B**). HaloBlast analyses of full length Risk1 and Risk2 proteins indicated distinct profiles, with Risk2 sharing low similarity to Legionellales kinases (**FIG. 5C**). Structural analyses corroborated this result, with limited regions of Risk1 and Risk2 modelling best to structures of *Helicobacter pylori* CtkA ^108^ and *L*. *pneumophila* LepB ^104^, respectively (**FIG. 5E,F**). Comparison of Risk1 and Risk2 PI3/4 active sites revealed 1) juxtapositioned aromatic residues in their catalytic loops, 2) the presence of a postively charged residue in the acivation loops of most Wortmannin-sensitive kinases (Risk1 and human class 1 and 2 PI3 kinases), and 3) greater sequence length between catalytic and activation loops in Risk2 proteins (**FIG. 5F**). Furthermore, only LepB and some rickettsial Risk2 proteins have the catalytic loop Asp replaced by Ser. These collective observations indicate two divergent PI3/4 kinases encoded in most rickettsial genomes (**FIG. 2**), leading us to posit that Risk2 is a PI4 kinase that complements the PI3 kinase activity of Risk1 ^56^.

We determined a remarkable connection between Risk2 and another *Rickettsia* effector, Sca4, which is highly conserved in *Rickettsia* species and implicated in intercellular spread by reducing mechanotransduction at cell-cell junctions ^76, 109^. The Sca4 C-terminal region has eukaryotic-like **vinculin binding sites** (**VBS**) that reduce vinculin-α- catenin interactions, which facilitates neighboring cell engulfment of *Rickettsia*-induced protrusions. The N-terminal region, shown by Schuenke and Walker ^110^ to elicit anti-rickettsia antibodies (Pfam: 120_Rick_ant), has a novel structure posited to guide assembly of the VBSs at host plasma membranes ^111^. Curiously, the plasmid pRPR of *R*. *peacockii* carries two adjacent genes encoding a divergent Risk2 and the N-terminal Sca4 domain ^112^ (**FIG. 5F**,**G**). This led to discovering a cohort of *Tisiphia* species and other MAGs encoding chimeric Pat2-Sca4-NT proteins, as well as numerous other protein architectures with the Sca4-NT domain (**FIG. 2**; **FIG. S5**; **FIG. 5G**). We renamed this pervasive domain **Schuenke Walker Antigen** (**SWA**) and suggest using **SWA**- **modular protein** (**SWAMP**) or coupling “SWA” with another domain when function is apparent; e.g., SWA-PIK (for PI kinase) or SWA-VBD (vinculin-binding domain). In accord with prior observations on high rates of recombination shaping Sca architectures ^77, 113, 114^, these results indicate the SWA domain is a recurrent module that may shuttle various types of effector functions to common host cell targets, particularly membranes.

#### From many forms, one descendent

Early in host infection, pathogens *R*. *typhi* and *R*. *rickettsii* secrete the REM RARP-2, which traffics to the endoplasmic reticulum and Golgi apparatus, leading to **trans-Golgi network** (**TGN**) fragmentation and ultimately perturbed protein transport to the host cell surface ^57, 115^. Like RalF, RARP-2 has a C-terminal tail that binds RvhD4 (**FIG. 1B**); furthermore, the protein has well delineated N-terminal protease and C-terminal ANK repeat domains (**FIG. 6A**). The protease domain has minimal analogy to clan CD protease families (C13 legumain, C14 caspase 1, C11 clostripain, and C25 gingipain R), which share a common fold that arranges a His and Cys catalytic dyad ^116^. This active site is essential for RARP-2 fragmentation of TGN ^115^ and also contributes to the lytic plaque phenotype of virulent *R*. *rickettsii* strains ^57^. The ANK repeat domain is atypical among most ANK repeat-containing proteins ^117^, as the composition of each repeat is highly similar in length and identity (**FIG. 6B**; **FIG. S6A-C**) despite a highly variable repeat number across orthologs, even at the strain level in most cases (**FIG. 6C**). RARP-2 active site mutants still traffic to perinuclear membranes, indicating that the ANK domain drives subcellular localization. However, shorter repeats (4 in attenuated strains Iowa *R*. *rickettsii*) do not disrupt TGN fragmentation relative to those of pathogenic *R*. *rickettsii* ^115^, suggesting a larger ANK domain is required for proper localization.

**FIGURE 6.**
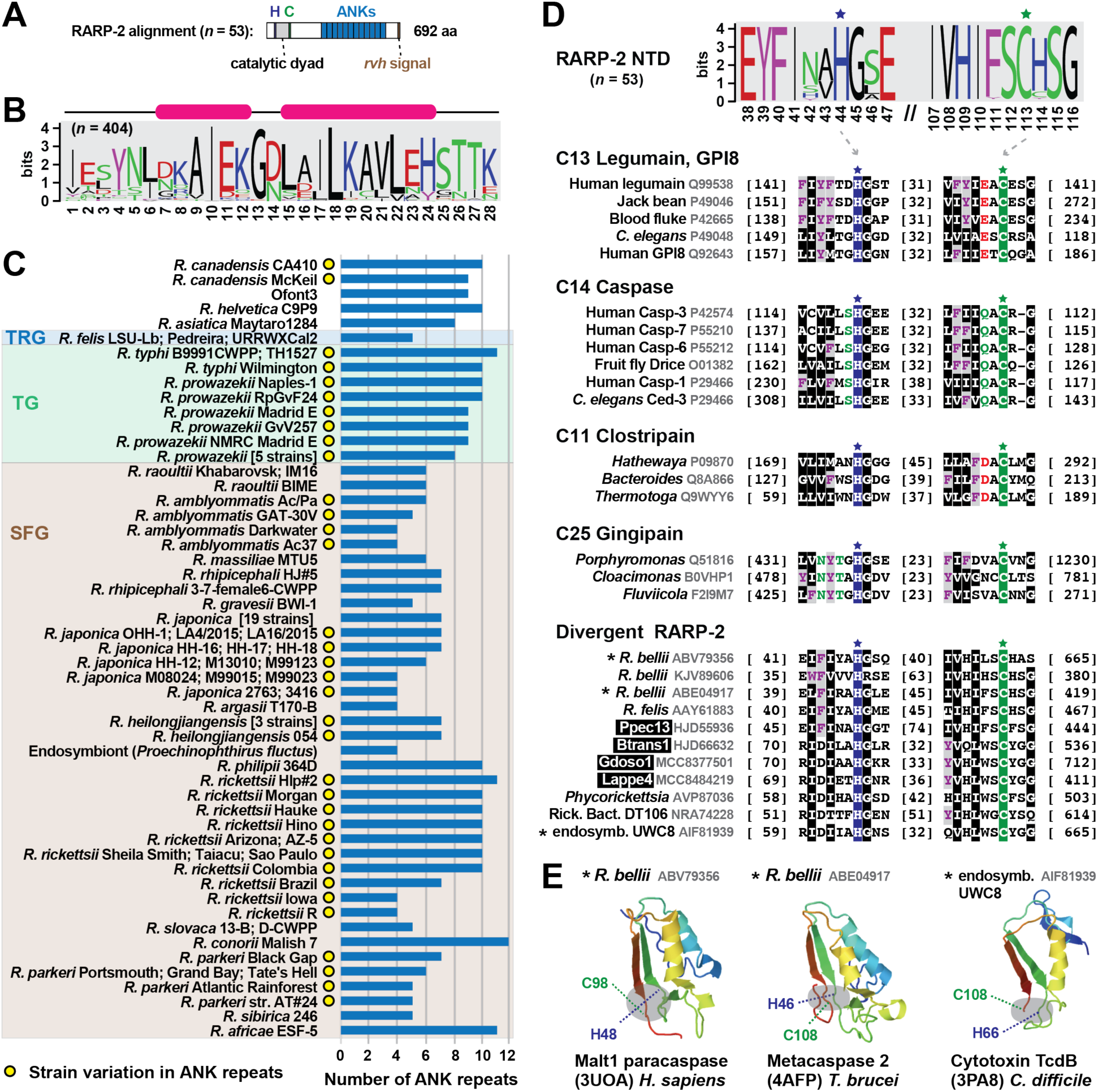
RARP-2 architecture is derived from multiple divergent forms. Black boxes provide short names for MAGs from Davison *et al*. ^72^. Amino acid coloring is described in the **FIG. 3** legend. Sequence logos constructed with WebLogo 3 ^187^. Sequence information in **Table S2**. **(A)** General architecture of RARP-2 proteins deduced from an alignment of 53 non-redundant RARP-2 proteins using MUSCLE ^186^ (default parameters). **(B)** Consensus sequence for the RARP-2 ANK repeat deduced from 404 repeats. **(C)** Depiction of the 53 non-redundant RARP-2 proteins with ANK domain repeat number provided. For brevity, some strain names are not shown for *R. prowazekii*: Chernikova, Katsinyian, Dachau, BuV67-CWPP, Rp22; *R. japonica*: YH, DT-1, HH-1, HH06154, HH07124, HH07167, MZ08014, Nakase, PO-1, Tsuneishi, HH-13, HH06116, HH06125, LON-151, M11012, M14012, M14024, SR1567, YH_M; *R. heilongjiangensis*: HCN-13; Sendai-29; Sendai-58. **(D)** RARP-2 and dRARP-2 proteins possess N-terminal domain clan CD cysteine protease-like active sites ^116^. Sequences were manually aligned to illustrate the conservation across all diverse protein groups. ‘Rick. endo. UWC8’, endosymbiont of *Acanthamoeba* str. UWC8 (not shown in FIG. 1C but closely related to endosymbiont of *Acanthamoeba* str. UWC36 in the Midichloriaceae). **(E)** Insight on RARP-2/dRARP-2 structure. Asterisks indicate proteins from panel **D** that were used in Phyre2 ^192^ searches to identify template structures for modeling ^118–120^. A complete structure of *R*. *typhi* RARP-2 predicted with Alphafold ^194, 195^ corroborates these dRARP-2 models and indicates deviations on a common effector architecture (FIG. S6C).

Probing recently sequenced genomes and MAGs did not reveal RARP-2 sequences in the **Bellii Group** (**BG**) rickettsiae or other Rickettsiaceae genomes (**FIG. 2**), consistent with prior observations that RARP-2 is unique to derived *Rickettsia* lineages ^88^. Yet, by focusing on the N-terminal protease domain, we discovered 56 **divergent RARP-2** (**dRARP-2**) proteins that possess the clan CD active site architecture (**FIG. 6D**). These proteins were binned into six groups (**FIG. S6D**) that have very different ANK repeat domain identities (data not shown); further, several could be modeled to structures of eukaryotic ^118, 119^ and prokaryotic ^120^ clan CD members (**FIG. 6E**). dRARP-2 proteins are predominantly found in BG rickettsiae and *Tisiphia* genomes, but likely shuttle in the intracellular mobilome given that one is carried by a Midichloriaceae species (endosymbiont of *Acanthamoeba* str. UWC8 ^121^). Based on the discordant genomic distribution of RARP-2 and dRARP-2 (**FIG. 1C**; **FIG. 2**) and the strong bias of RARP-2 in vertebrate-associated species, we speculate that RARP-2 and dRARP-2 may be tailored for similar functions related to TGN fragmentation yet well diverged to allow recognition of targets specific to disparate eukaryotic hosts. This is reminiscent of the recent discovery that *R*. *parkeri* utilizes different factors for apoptosis induction in ticks versus mammals ^122^.

#### cREMs with Unknown Function

For five *R*. *typhi* **hypothetical proteins** (**HPs**) previously shown to interact with RvhD4 (cREM-1-5; **FIG. 1B**), MAG analyses provided substantial clarity on the mechanisms of evolution shaping their architectures. Four of these proteins are described below in light of newfound gene fission/fusion and duplication events (cREM-1, cREM-2, and cREM-4), as well as a greater role of conjugative transposons shaping *Rickettsia* evolution (cREM-5). Unexpectedly, the small cREM-3 (∼93 aa) was determined to have widespread conservation in Rickettsiales yet also exist in certain other *Proteobacteria* (**FIG. 1C**; **FIG. 2**). While likely not a REM, our analyses revealed a potential structure associated with this curious protein (**FIG. S7C**).

#### Cryptic gene fission and duplication obscured by rapid divergence

For cREM-1 and cREM-2, we utilized phylogeny estimation in conjunction with sequence analysis to predict gene fission (cREM-1) and duplication (cREM-2) events behind the evolution of these proteins. Neither protein was found outside of Rickettsiaceae (**FIG. 1C**; **FIG. 2**). cREM-1 proteins are streamlined from larger modular *Tisiphia* proteins that harbor the entire cREM-1 sequence as a domain; accordingly, we named these **divergent cREM-1** (**cREM-1d**) (**FIG. 7A**). Some rickettsiae carry cREM-1 tandem duplications, though most genomes have one conserved gene and the second pseudogenized (**FIG. S7B**; red clade). Curiously, cREM-1 proteins have similarity to a repeat region within the *O*. *tsutsugamushi* effector OtDUB (**FIG. 7A**; **FIG. S7A**). This region in OtDUB binds clathrin adaptor-protein complexes AP-1 and AP-2 and harbors a cryptic Rac1 GEF domain ^123–125^. This indicates cREM-1 proteins may have evolved from larger modular proteins with functions tailored to the eukaryotic cytosol, with repeat regions of these large effectors streamlining to smaller proteins encoded by tandem genes (**FIG. 7B**; **FIG. S7A**).

**FIGURE 7.**
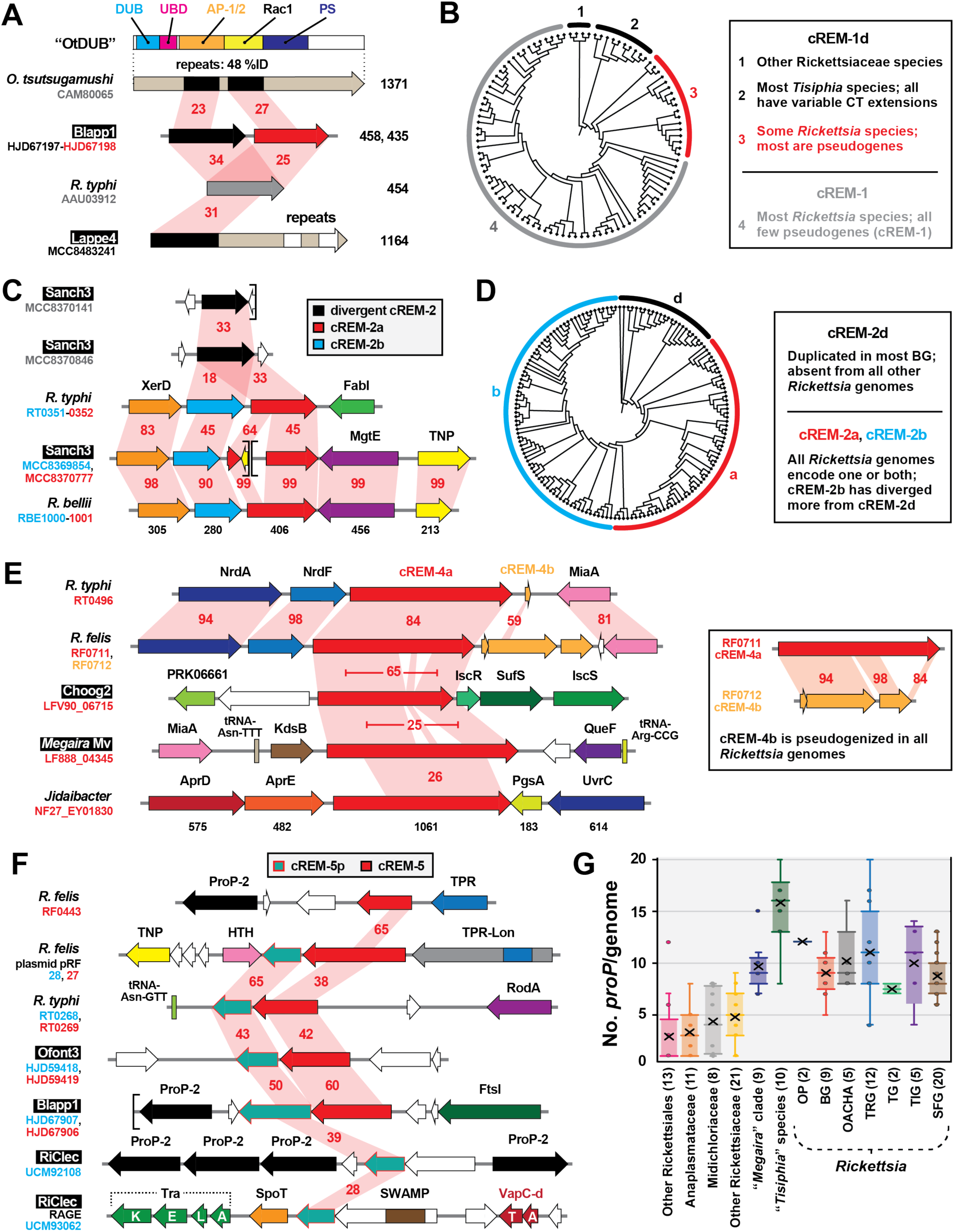
Gene fission and duplication has shaped the architectures of four candidate REMs. Black boxes provide short names for 29 MAGs from Davison *et al*. ^72^. Gene regions drawn to scale using PATRIC compare region viewer tool ^198^. Sequence information is provided in **Table S2**. **(A)** Similarity between *O*. *tsutsugamushi* effector OtDUB (CAM80065), divergent cREM-1 (cREM-1d), and cREM-1 proteins. OtDUB characterized domains: deubiquitinase (light blue), ubiquitin-binding (pink), cryptic Rac 1-like guanine nucleotide exchange factor (yellow), clathrin adaptor-protein complexes AP-1 and AP-2 (orange), phosphatidylserine-binding (gray) ^5, 123, 201^. Red shading and numbers indicate % aa identity across pairwise alignments. **(B)** Cladogram depicting phylogeny estimation of 102 cREM-1d and cREM-1 proteins (see FIG. S7A for phylogram and methods). Inset describes classification of cREM-1 proteins, with clade colors matching the protein colors in the schema in panel **A**. **(C)** Diversification of cREM-2 proteins via duplication. **(D)** Cladogram depicting phylogeny estimation of 158 cREM-2d, cREM-2a, and cREM-2b proteins (see FIG. S7B for phylogram and methods). Inset describes classification of cREM-2 proteins, with clade colors matching the protein colors in the schema in panel **C**. **(E)** Ancient gene duplication of cREM-4 and location of cREM-4 genes in select Rickettsiales species. The cREM-4 pentapeptide repeat domain is illustrated in FIG. S7D. **(F)** cREM-5/5p loci occur in variable genomic regions, including plasmids and RAGEs. The complete RAGE for RiCle is illustrated in FIG. S7F. **(G)** Genomic distribution of ProP genes in Rickettsiales genomes. OP, *Rickettsia* endosymbionts of *Oxypoda opaca* (Oopac6) and *Pyrocoelia pectoralis* (Ppec13); OACHA, *Rickettsia* endosymbionts of *Omalisus fontisbellaquei* (Ofont3) and *Adalia bipunctata*, *R*. *canadensis*, *R*. *helvetica*, and *R*. *asiatica*.

cREM-2 proteins belong to pfam17422 (DUF5410: specific to *Rickettsia* species). Our analyses identified a second DUF5410-like protein encoded adjacent to cREM-2 proteins in many *Rickettsia* genomes (**FIG. 2**). Neither of these tandem duplicates (designated cREM-2a and cREM2-b) contain Sec signal sequences or other predictable features (**FIG. 7C**). Further, some BG rickettsiae and *Tisiphia* species harbor a third **divergent cREM-2** (**cREM-2d**) that is absent from derived *Rickettsia* lineages. With the assumption that cREM-2d are an ancestral form, phylogeny estimation indicates cREM-2b proteins are more divergent than cREM-2a proteins (**FIG. 7D**), though all three protein architectures share high conservation in dozens of residues within the central region of these proteins (**FIG. S7B**).

cREM-4 proteins also show evidence of an ancestral duplication (**FIG. 7E**), though no genomes contain a complete duplicate gene, indicating a consistent pseudogenization event that rapidly followed cREM-4 duplication (**FIG. 2**). Despite their large size (∼950 aa) these proteins contain only one observable feature, a small internal **pentapeptide repeat** (**PR**). While widespread in diverse bacterial proteins, PR function is generally unknown, though some bacterial PR-containing proteins can interact with DNA binding proteins ^126^ and contribute to virulence ^127^ (**FIG. S7D**). cREM-4 proteins are encoded in certain other Rickettsiaceae genomes, and like dRARP-2, a single Midichloriaceae species (“*Candidatus* Jidaibacter acanthamoeba” ^30^) encodes a cREM-4 protein. While cREM-4 of BG rickettsiae lack the PR (**FIG. S7D**), nearly all derived *Rickettsia* genomes encode a complete cREM-4 protein, indicating retention of a conserved function after an ancestral duplication.

#### LGT of cREM-5 as a two-gene module across select species

cREM-5 proteins were previously noted for their restricted distribution in TG rickettsiae and *R*. *felis* (both in the chromosome and on plasmid pRF) ^128^. Our analyses yielded several novel findings. First, while absent from any **Spotted Fever Group** (**SFG**) or **Tamurae/Ixodes Group** (**TIG**) rickettsiae, cREM-5 proteins are highly conserved in all BG rickettsiae genomes, as well as in a few *Tisiphia* genomes (**FIG. 2**). Second, most cREM-5 genes have an associated protein, **cREM-5 partner** (**cREM-5p**), encoded immediately downstream (**FIG. 7F**). Despite conserved regions (**FIG. S7E**), neither protein has detectable domains or similarity to proteins in other Rickettsiales (**FIG. 2**). Third, cREM-5/5p genes have a strong co-occurrence with PropP-2 genes (black, **FIG. 7F**). ProP (Proline Betain Transporters of the Major Facilitator Superfamily) function in osmoregulation ^129, 130^ are proliferated in *Rickettsia* genomes, with seven conserved groups (PropP1-7) containing species specific duplications ^90, 131^. Why specifically PropP-2 genes cluster near certain cREM-5/5p loci is unclear, though insight from MAGs illuminated a previously unrealized point in Rickettsiales evolution where ProP proliferation occurred (**FIG. 7G**).

Finally, cREM-5 modules are found in recombination hotspots and other less conserved genomic regions, indicating LGT behind their evolution in *Rickettsia* genomes. Aside from cREM-5/5p on plasmid pRF, one copy of cREM-5p from the RiClec (Endosymbiont of *Cimex lectularius*) genome is found on a conjugative transposon termed **Rickettsiales Amplified Genetic Elements** (**RAGE**) (**FIG. 7F**). RAGE are integrative and conjugative elements present on certain *Rickettsia* plasmids and chromosomes ^90, 91, 132^, as well as proliferated and scattershot in *O*. *tsutsugamushi* genomes ^133, 134^. Cargo genes, or those piggybacking on RAGEs at indiscriminate insertion sites, have functions mostly related to the stringent response and metabolism, defense and resistance, and adaptation to host cells (e.g., ProP genes are shuttled by RAGE). The addition of cREM-5p and SWAMPs, as well as a myriad of TA modules, to the list of RAGE cargo (see **FIG. S7F**) indicates these mobile elements play a role in disseminating pathogenicity factors that was previously unappreciated.

### cREMs with Characterized Function

Several *Rickettsia* proteins that lack N-terminal Sec signals have either been well characterized for their roles in subverting host cell processes (e.g., Sca4 and RickA) or possess features that implicate them in targeting host molecules (e.g., VapC and other toxins within TA modules). Until secretory pathways for these molecules are characterized, we consider them cREMs (**FIG. 1B**). MAG analyses of these proteins have generated novel insight on the structure and evolution of domains targeting the host actin cytoskeleton. Furthermore, a greater appreciation for toxin architecture and distribution indicates the *rvh* T4SS may still function in congener killing despite the host-dependent lifestyle of the derived Rickettsiales species.

#### Insight on Rickettsia interactions with host actin

With their RalF proteins lacking VPRs and their SWAMPs lacking VBDs (Sca4), *Tisiphia* species may interact with host cell actin cytoskeleton differently than *Rickettsia* species (**FIG. 8A**). We analyzed another host actin-associated protein, RickA, which some *Rickettsia* species use for intracellular motility and possibly intercellular spread ^135–138^. While no association with the *rvh* T4SS has been characterized, RickA localizes to the bacterial surface in the absence of a Sec signal peptide ^138^. RickA directly activates host Arp2/3 complexes through an architecture that mimics host **nucleation promoting factors** (**NPF**) ^78, 139^. Surprisingly, we discovered several RickA proteins from *Tisiphia* MAGs that differ in their C-terminal architectures relative to SFG rickettsiae RickA proteins characterized in actin polymerization (**FIG. 8B**; **FIG. S8A**). While the functional relevance of these differences is unclear, we gained further insight on the N-terminal structure of all RickA proteins. The substantial increase in diversity from MAGs illuminated a large (∼95 aa) repeat region enclosing the G actin-binding domain, with each repeat highly conserved in hydrophobicity and predicted structure (**FIG. 8B**; **FIG. S8B**). We envisage that this conserved region may facilitate docking of G actin to the G actin-binding domain and overall positioning of the N-terminus to **Wiskott-Aldrich syndrome protein homology 2** (**WH2**) motifs at the C-terminal region.

**FIGURE 8.**
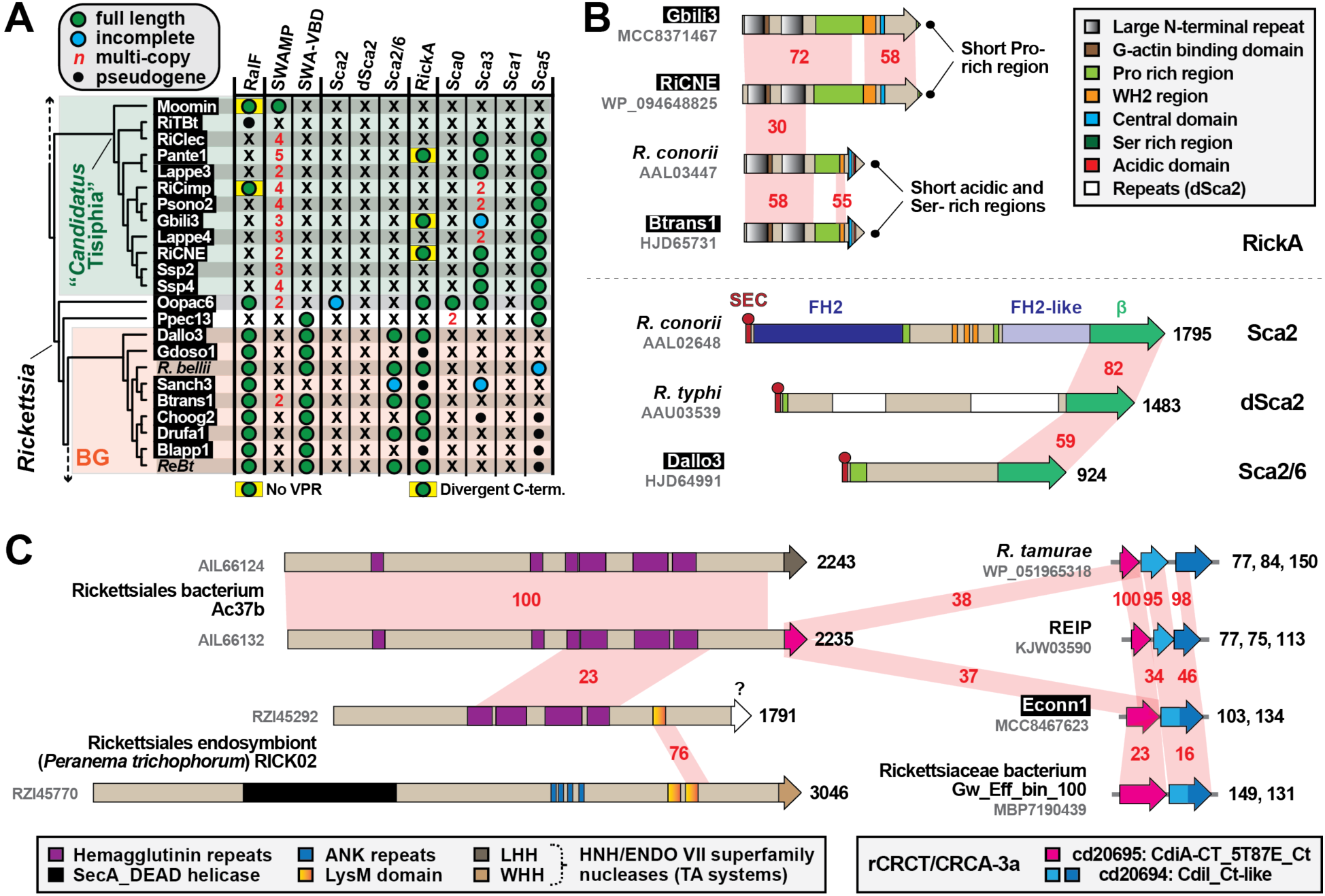
MAG analyses lend insight on *Rickettsia* interactions with host actin cytoskeleton and *rvh* T4SS function. Black boxes provide short names for 29 MAGs from Davison *et al*. ^72^. **(A)** MAGs shed light on the evolution of *Rickettsia* factors behind host actin polymerization and invasion. *Tisiphia* and BG rickettsiae taxa, as well as SWAMP, SWA-VBD, RickA, and RalF info, is from FIG 2. The passenger domains of *R*. *conorii* Sca2 (AAL02648), *R*. *typhi* dSca2 (AAU03539), *R*. *bellii* Sca2-6 (ABE05361), *R*. *conorii* Sca0 (AAL03811), *R*. *typhi* Sca3 (AAU03915), *R*. *typhi* Sca1 (AAU03504) and *R*. *typhi* Sca5 (AAU04158) were used in BlastP searches directly against *Tisiphia* and BG rickettsiae genomes. Passenger domains and linker sequences were delineated as previously shown ^88^. *R*eBt, *Rickettsia* spp. MEAM1, wb, and wq (endosymbionts of *Bemisia tabaci*). **(B)** Some *Rickettsia* genomes encode one or more host actin nucleation proteins. ***Top***: *Tisiphia* and *Rickettsia* RickA proteins share a large N-terminal repeat domain but diverge at their C-termini. WH2, Wiskott-Aldrich syndrome protein homology 2 domain. Further details on RickA architecture are provided in FIG. S8. ***Bottom***: Sca2, d-Sca2 and Sca2-6 proteins have a common autotransporter domain (β) but divergent passenger domains. FH2, formin homology 2. Sca2 mimics host formin actin nucleators ^143^ to recruit and polymerize actin for intracellular motility and intercellular spread ^141, 142^. The functions of dSca2 and Sca2/6 are unknown. **(C)** The mobile nature of CDI-like/Rhs-like C-terminal toxin/antidote (CRCT/CRCA) modules across diverse rickettsial genomes. Schema shows integration of CRCT/CRCA modules into larger polymorphic toxins (hemagluttinin-like toxins, LysM-like peptidoglycan/chitin-targeting toxins, etc.), as well as CRCT/CRCA modules independent of larger toxins. The toxin warhead for RZI45292 is unknown. Further details are provided in FIG. S9.

Despite activation by RickA, the specific role of host Arp2/3 complexes during rickettsial infection is unclear, perhaps due to different species utilized across studies garnering contrasting results ^137, 138, 140^. Specifically, actin-based motility in certain *Rickettsia* species is carried out by a second NPF, Sca2, that polymerizes actin independent of Arp2/3 complexes ^141, 142^. The passenger domains of Sca2 mimic eukaryotic formins by elaborating ring-like structures to elongate actin, with intervening Pro-rich regions and WH2 domains incorporating profilin-actin for elongation and recruiting actin monomers for nucleation, respectively ^143^. Thus, at least for species carrying both RickA and Sca2 (most SFG rickettsiae), RickA-mediated Arp2/3 activation may play a greater role early in infection, possibly in filopodia formation during invasion ^79, 142, 144, 145^. Still, few *Rickettsia* species outside of SFG rickettsiae encode Sca2 proteins with intact formin-like passenger domains (**FIG. 8B**) and some of these species lack RickA genes as well ^90, 112, 146^ (**FIG. 8A**; **FIG. 2**). This implies that host actin polymerization for motility is an expendable trait for most *Rickettsia* species, and that Arp2/3 recruitment and activation during invasion can be instigated by other bacteria-driven processes, i.e. Arf recruitment to the plasma membrane for inducing PI shifts required for filopodia formation ^147^.

MAG analyses indicate that, barring acquisitions via LGT, RickA and RalF were likely present before the diversification of major *Rickettsia* lineages, whereas Sca2 appeared later in *Rickettsia* evolution (**FIG. 8A**). Further, we polled MAGs for the presence of genes encoding the four major autotransporters (Sca0, Sca1, Sca3, and Sca5) with known (or anticipated) functions in host cell binding and/or invasion ^148–152^. Remarkably, Sca3 is predominant in *Tisiphia* genomes despite a very limited distribution in *Rickettsia* species (restricted mostly to TG and TRG rickettsiae ^88^). Further, BG rickettsiae are counter to most other rickettsiae in lacking both Sca0 and Sca5, the dominant proteins of the characterized *Rickettsia* S-layer ^153^. Collectively, these analyses show that *Rickettsia* factors described in host cell invasion and actin cytoskeleton subversion are sporadically encoded across genomes, indicating host specificity and/or expendability in their contribution to the intracellular lifestyle.

#### A repurposed or multi-purposed rvh T4SS?

Aside from secreting effectors that target host cellular processes, evidence is mounting for intracellular bacteria utilizing large **contact dependent growth inhibition** (**CDI**) and **Recombination hotspot** (**Rhs**) toxins for interbacterial antagonism ^154, 155^. We recently identified a few rickettsial genomes encoding specialized TA modules that some bacteria integrate into CDI and Rhs toxins to expand toxic activities ^156–159^. Widespread in bacteria, these **CDI-like/Rhs-like C-terminal toxin and antidote** (**CRCT/A**) modules are extremely polymorphic, variable at the species- and strain-level, and found either associated with larger toxins or alone as small TA modules ^160^. The two types of ***Rickettsia* CRCT/A** (**rCRCT/A**) modules we identified, rCRCT/A-1 and rCRCT/A-3a, were once associated with large Rhs-like toxins that have mostly degraded ^73^. rCRCT/A-1 modules are highly divergent from other characterized CRCT/A modules and predominantly occur in Actinomycetia and Cyanobacteria genomes. Only two *Rickettsia* species, *R*. *tamurae* and *R*. *buchneri*, harbor rCRCT/A-1 modules; however, the “*Cand.* J. acanthamoeba” genome encodes one as an independent module and one integrated into a large modular hemagglutinin toxin with nuclease and peptidase domains. MAG analyses herein discovered several more rCRCT/A-1 modules mostly in *Tisiphia* genomes associated with pseudogenized hemagglutinin-like toxins (**FIG. 1C**; **FIG. 2**).

rCRCT/A-3a modules resemble the prototype CDI TA module (CdiA-CT/CdiI), wherein the nuclease CdiA-CT targets tRNAs in recipient cytosol ^156^. CdiA-CT/CdiI is associated with a large modular protein (CdiA) that joins with a second protein (CdiB) as a **type Vb secretion system** (**T5bSS**) to deliver the toxin into neighboring bacteria ^158, 160, 161^. However, rCRCT/A-3a modules (and all Rickettsiales genomes) lack CdiB genes. This type of CRCT/A module is widespread in proteobacterial genomes ^158^. MAG analyses also revealed more rCRCT/A-3a modules in *Rickettsia* genomes and a much higher presence of single antidotes (**FIG. 1C**; **FIG. 2**), possibly indicating greater selection for defense against toxins versus toxin secretion. Additionally, a rCRCT/A-3a module was found integrated into a large hemagglutinin-like toxin in the Rickettsiales endosymbiont Ac37b, an early-branching Rickettsiaceae species that can co-infect ticks with SFG rickettsiae (**FIG. 8B**; **FIG. S9**). Remarkably, this species also carries an identical hemagglutinin-like toxin but with a divergent warhead of the HNH/ENDO VII nuclease superfamily, illustrating the integrative nature of diverse CRCT/A modules. Further, two toxins in the genome of the Rickettsiales endosymbiont of *Peranema trichophorum* (Midichloriaceae) carry different C-terminal toxins, as well as lysin motifs (LysM) that often occur in cell wall degrading enzymes ^162^. Collectively, our analyses illuminate diverse CRCT/A modules in the Rickettsiales mobilome that equip bacteria with weapons for interbacterial antagonism.

Rickettsiales species may also utilize **filamentation induced by cAMP** (**FIC**) proteins and type II TA modules for interbacterial antagonism. Some intracellular bacteria harbor FIC domain-containing proteins ^155^ and several human pathogens secrete effectors with FIC domains into host cells to subvert cellular processes ^163^. Furthermore, a recent report illustrated that *Yersinia pseudotuberculosis* utilizes a FIC domain effector, CccR, that alters conspecific gene expression and inhibits congener growth ^164^. Many *Rickettsia* genomes encode multiple divergent FIC proteins (**TABLE S2**); however, to our knowledge, none of these proteins are known to be secreted by rickettsiae. Similarly, *Rickettsia* species also harbor a myriad of diverse type II TA modules, with many found on RAGE (e.g., **FIG. S7F**; **FIG. S7F**) or plasmids ^91, 165^. Only one module, VapBC of *R*. *felis*, has been characterized. Structural analysis revealed the nature of antidote (VapB) binding to toxin (VapC) ^166^, and VapC possesses toxic RNase activity when expressed in bacterial or eukaryotic host cells. We previously showed that *Rickettsia* genomes encode VapBC and/or a divergent module (VapBC-d) ^88^, and MAG analysis confirmed this observation (**FIG. 2**). Further, in light of the new genomic diversity, more discrete VapBC loci are encoded in many genomes (data not shown), as well as other type II TA modules (e.g., those encoding ParE, BrnT, and RatA toxins) that have yet been characterized (**TABLE S2**).

Finally, MAG analyses doubled the number of Rickettsiaceae proteins harboring domains found in RP toxins, particularly those of *Wolbachia* cytoplasmic incompatibility inducing nucleases (CinB) and deubiquitinases (CidB), as well as the *Spiroplasma* male killer toxin deubiquitinase (Spaid) ^167–171^ (**FIG. S10**). Many of these toxins are substantially large and modular, encoding numerous domains with uncharacterized effects on host cells ^5, 172, 173^ (e.g., see **FIG. 3D**). The increasing number of RP toxins (and antidotes when present) in rickettsiae, particularly in species associated with male-killing and parthenogenesis phenotypes, indicates undiscovered molecular mechanisms underpinning these modes of RP. Like the rCRCT/A modules, FIC toxins, and type II TA modules, these RP toxins all lack characterized secretion pathways. Thus, the *rvh* T4SS cannot be ruled out as a secretion pathway for any of these potential effectors, though a T1SS conserved in all Rickettsiales ^88^ (and possibly other unappreciated routes) could be involved.

### Power and Efficacy of MAG Diversity

The inclusion of diverse MAGs in the assessment of *rvh* effector evolution has provided several key insights. First, like the *rvh* T4SS ^41^, many REMs and cREMs contain gene duplication. However, unlike the *rvh* machine, effector duplication seems to define basal lineages (*Tisiphia*, BG rickettsiae, and other Rickettsiaceae) and tends to lead to retention of only one protein in the derived *Rickettsia* groups. Still, divergent forms arising from duplication stand to inform on effector function, particularly if derived proteins are utilized for vertebrate cell infection.

Second, the sparse distribution of polled effectors outside of *Rickettsia* genomes indicates they originated after the divergence of rickettsial families. In some cases, analyses strongly implicate LGT for acquisition of effectors, with a particular bias from Legionellales (e.g., RalF, patatins and PIKs) and other aquatic microbes. This supports the “intracellular arena” hypothesis for the gain of similar effectors in divergent pathogens that occupy common hosts (i.e. protists and arthropods) ^174^. It also corroborates our earlier observations that LGT, particular by RAGEs and plasmids, offsets reductive genome evolution in rickettsiae ^26, 90^. A more recent study reached a similar conclusion via discovery for gene gain shaping Chlamydiae genome architecture, despite the reduced size of most chlamydial genomes ^175^. MAGs have also provided a greater appreciation for Legionellales diversity and revealed that the major host-adaptive features (i.e. the *dot*/*icm* T4SS and a few conserved effectors) were established in the last common Legionellales ancestor ^176^. This is consistent with the discovery by Schön *et al*. of the *rvh* T4SS in ancestral Rickettsiales ^28^; however, we caution that strict *rvh* repurposing from congener killing to host parasitism is premature until the secretory pathways of the numerous toxins described above are experimentally determined.

Third, MAGs help bridge the gap between research on microbial ecology and human pathogenesis, revealing genome evolutionary and architectural traits that are underappreciated due to biases of clinical isolates or more common environmental strains on public databases ^71, 72^. Our discovery here of REMs and cREMS on novel plasmids and RAGEs accentuates this point, indicating that such genetic elements may be underestimated for roles in rickettsial biology due to the strong bias of high passage clinical isolates on databases. This is particularly relevant in light of the recent demonstration that the ***Rickettsia* regulator of actin motility** (***roaM***) is often pseudogenized in highly passaged laboratory strains, suggesting serial passage in cell culture can eliminate essential genes lacking environmental selective pressure (in this case the arthropod cytosol) ^177^.

Fourth, the most profound insight gained from our work shows how MAG analyses often illuminate novel architectures for well-studied virulence factors. Unearthing new effector designs provides clues on how general foundations are tailored to different hosts and host cell processes. This is epitomized by our discovery of novel RalF-like proteins with SCDs substituted for ARF-interacting domains, which not only fortifies the literature on *Legionella* and *Rickettsia* RalF-mediated host ARF recruitment ^55, 81, 84, 86, 87^, but also pinpoints the rise of the actin-targeting VPR regions in *Rickettsia* RalF proteins after the divergence from *Tisiphia* species. Combined with numerous other novel effector architectures identified herein, this highlights remarkable recapitulation on mechanisms for mimicking eukaryotic functions that exist beyond *Rickettsia* and other human pathogens and are widespread in the environment. We assert that widening the comparative genomics lens will allow evolution, which has already matched effector form and function to host environments, to guide experimental designs and reinvigorate pathogen effector research.

Finally, as the landscape of *Rickettsia* pathogenesis undergoes gradual change due to virulence factor characterization and immunological studies ^1, 2^, the traditional designation of SFG and TG rickettsiae as the major lineages defining the genus has become grossly outdated. A substantial spike in TRG rickettsiae diversity ^72, 178, 179^, coupled with robust genome-based phylogeny estimations and phylogenomic analyses ^28, 72^, make the common ancestry of TG and TRG rickettsiae incontrovertible. Prior bias in genome sequences for SFG rickettsiae portrayed TG rickettsiae as unique by smaller genome size and greater pseudogenization relative to all other rickettsiae. However, our focus on *rvh* effectors across a highly diverse and unbiased genomic sampling shows that all of the major *Rickettsia* groups (BG, TRG, TG, TIG, and SFG rickettsiae) have distinct evolutionary trajectories of gene gain, loss, and modification (**FIG. 2**). Thus, MAGs have exposed far greater *Rickettsia* diversity than previously realized, though long ago conjectured by environmental sampling ^180^. These data, as well as careful dissection of the attributes distinguishing *Tisiphia* and *Rickettsia* species, will be paramount for deciphering how human pathogens have emerged, possibly multiple times, from this veritable bevy of endosymbiont diversity. Furthermore, an understanding of environmental genomic richness, particular in mobile element diversity, may help forecast the next serious rickettsial diseases to emerge.

## CONCLUSION

Discovery and analyses of MAGs has greatly impacted the landscape of Rickettsiology, adding substantial diversity and dispelling the long-held dogma for an ancestral link to the mitochondrial ancestor. Despite predicted extracellular lifestyles, basal rickettsial species carry the *rvh* T4SS and likely use it as a congener killing machine. Our study coupled a robust evolutionary framework with the inspection of over two dozen known or predicted *Rickettsia rvh* effectors to provide insight on the origin of mechanisms for host cell subversion and obligate intracellular parasitism. Though focused in taxonomic scope, this experimental design is amenable to probe the origins of virulence factors in any human pathogen with representation in the diverse treasure trove of MAG data. At bare minimum, our work demonstrates that utilizing MAGs in comparative approaches greatly enlightens dialogue on mechanisms of pathogenesis.

## MATERIALS AND METHODS

### Rickettsiales Phylogeny Estimation

Robust genome-based phylogeny estimations for Rickettsiales ^28^ and *Rickettsia*-*Tisiphia* ^72^ were used as benchmarks to evaluate our estimated phylogenies based on single or concatenated *rvh* proteins. We polled the rich MAG diversity on the NCBI database for the presence of *vir*-like T4SS genes possessing *rvh* hallmarks ^37, 41, 44^ (i.e., RvhB8, RvhB9 and RvhB4 duplication, multicopy RvhB6, no VirB5 analog; **FIG. S1**). Provided that many MAGs and certain genome assemblies are likely incomplete, we limited our dataset to assemblies containing both RvhB4-I and RvhB-II, except for a few cases where strong evidence from other *rvh* genes indicated a Rickettsiales assembly. A total of 153 genome assemblies were retained for further analyses: 1) 93 Rickettsiaceae genome assemblies (including the 28 MAGs from Davison *et al*. ^72^ and another 15 previously unanalyzed MAGs), 2) 14 and 9 genome assemblies from Anaplasmataceae and Midichloriaceae, respectively, 3) the “*Candidatus* Deianiraea vastatrix” (Deianiraeaceae) genome assembly, and 4) 33 environmental MAGs likely comprising Deianiraeaceae, Athabascaceae or Mitibacteraceae (nine previously analyzed by Schön et al. ^28^) (**Table S1**; **FIG. S2**).

Only RvhB4-I and RvhB4-II proteins were included in phylogeny estimation, as alignments of other Rvh proteins were extraordinarily variable across the selected taxa (data not shown). RvhB4-I and RvhB4-II proteins were separately aligned using MUSCLE (default parameters). Each alignment included *Agrobacterium tumefaciens* str. F4 VirB4, which was used as an outgroup to root estimated trees. RvhB4-I and RvhB4-II protein alignments were subsequently concatenated (1974 total positions, “unmasked alignment”). TRIMAL ^181^ was used to create a second alignment with less conserved positions masked (1613 total positions, “masked alignment”).

For both unmasked and masked alignments, a maximum likelihood-based phylogeny was estimated with PhyML ^182^, using the Smart Model Selection ^183^ tool to determine the best substitution matrix and model for rates across aa sites (LG (G+I+F) for both alignments). Branch support was assessed with 1,000 pseudo-replications. Trees were drawn using FigTree (https://github.com/rambaut/figtree/) and manually modified using Adobe Illustrator ™. Final trees were manually adjusted to place “*Candidatus* Sneabacter namystus” (which lacks the *rvh* T4SS) in a position on the phylogram suggested by prior phylogeny estimation ^184, 185^. All terminal taxa were assigned names based on NCBI database taxonomy (as of Feb. 26^th^, 2023), with some “short-names” taken from Davison *et al*. ^72^ (these are provided in black boxes throughout the figures). Rickettsial classification scheme (STG, Scrub Typhus Group; BG, Bellii Group; TRG, Transitional Group; TG, Typhus Group; TIG, Tamurae-Ixodes Group; SFG, Spotted Fever Group) follows our prior reports ^73, 165^.

### Phylogenomics Analysis

The RvhB4-based estimated phylogeny was used as a scaffold to complete a distribution matrix for REMs and cREMs. It was not our goal to assess the relative completeness of each MAG included in the matrix, yet to only assess if MAGs and other genome assemblies possessing a *rvh* T4SSs also include counterparts (homologs or analogs) to *Rickettsia* REMs and cREMs. REM/cREM assignment is based on prior studies implicating their secretion and/or interaction with RvhD4 (by bacterial 2-hybrid and/or coimmunoprecipitation assays) or presence of motifs known to target either congener bacteria or eukaryotic molecules ^55–57, 73–80^. Analyses of some REMs and cREMs illuminated more complex gene structures (duplications, gene streamlining, and gene fusions) that prompted expansion of the total effector dataset. A total of 26 proteins were analyzed within the final phylogenomic framework (**Table S2**).

### In silico Protein Characterization

Analyses of each REM and cREM dataset contained discrete workflows tailored to the level of effector conservation in Rickettsiales (and in some cases other bacteria), prior studies that included bioinformatics analyses, and identification of gene duplication, streamlining, or gene fusion. All individual workflows are described in the pertinent figure legends and/or supplemental figure legends. Only general bioinformatics analyses are described below.

#### Dataset compilation

*Rickettsia* REMs and cREMs were used as queries in Blastp searches to compile and analyze diverse proteins harboring significant similarity across the entire lengths of the queries. Analyses utilized our HaloBlast method, which is a combinatorial Blastp-based approach originally designed to interrogate proteins for LGT^26^. HaloBlasting compiles Blastp subjects from restricted taxonomic searches that theoretically decrease in similarity by sampling lower levels of bacterial classification. A general search strategy for rickettsiae entails individual Blastp searches against six distinct taxonomic databases: 1) “Rickettsia” (NCBI taxid 780)”; 2) “Rickettsiales” (taxid: 766) excluding “Rickettsia”; 3) “Alphaproteobacteria” (taxid: 28211) excluding “Rickettsiales”; 4) “Proteobacteria” (taxid: 1224) excluding “Alphaproteobacteria”; 5) “Bacteria” (taxid: 2) excluding Proteobacteria”; and 6) “minus bacteria”. Data subsets were constructed strictly using NCBI taxid and following the NCBI taxid hierarchy to identify “daughter” taxonomic groups. Typically, 500 subjects (if available) are retained per search. All subjects from each search were separately ranked by *Sm* score (= *b* * *I* * *Q*, where *b* is the bitscore of the match, *I* is the percent identity, and *Q* is the percent length of the query that aligned), a comparative sequence similarity score designed to de-emphasize highly significant matches to short stretches of the query in favor of longer stretches of similarity ^26^. The “halos” or separate database searches are then compared to one another to determine the taxon with the strongest similarity to the query sequences. These analyses usually make LGT apparent when divergent datasets contain top-ranking proteins more similar to the *Rickettsia* queries than more closely related datasets.

#### Protein characterization

Select proteins or domains (again, context-dependent) are typically compiled and aligned with MUSCLE using default parameters ^186^. To identify conserved regions, alignments are then visualized as sequence logos using WebLogo^187^. Domain analyses are performed by cross-checking predictions from the NCBI **Conserved Domains Database** (**CDD**) and EMBL’s **Simple Modular Architecture Research Tool** (**SMART**) ^188^. In some cases, proteins were evaluated for N-terminal signal peptides ^189^ and transmembrane-spanning regions ^190^. Alignments shown in the figures and supplemental figures are manually assessed for conservation, typically considering 80% of a position conserved (alignment size-dependent), with amino acid coloring scheme and assignment as follows: black, hydrophobic (Ala, Val, Iso, Leu, Pro, Met, Gly); gray, less hydrophobic (can include a minority of Try, Phe, Tyr); red, negatively charged (Glu, Asp); green, hydrophilic (Cys, Asn, Gln, Ser, Thr); purple, aromatic (Try, Phe, Tyr); blue, positively charged (His, Lys, Arg). Individual protein schemas were generated using Illustrator of Biological Sequences ^191^ with manual adjustment in Adobe Illustrator ™.

Protein structures were predicted using the **Protein Homology/analogY Recognition Engine** V 2.0 (**Phyre2**) ^192^; in some cases, published structures were retrieved from the Protein Data Bank ^193^ and used in one-to-one threading mode with Phyre2. For some effectors, we also evaluated structures generated with Alphafold ^194, 195^. Finally, some regions of proteins were analyzed for predicted secondary structure using JPred ^196^.

Phylogenies were estimated for some REM and cREM datasets, which were compiled uniquely for each case and utilized HaloBlast to obtain non-rickettsial taxa (if available). Alignments were not masked since masking eliminated too many informative positions. Maximum likelihood-based phylogenies were estimated with PhyML ^182^, using the Smart Model Selection ^183^ tool to determine the best substitution matrix and model for rates across aa sites. Branch support was assessed with 1,000 pseudo-replications. Trees were visualized and drawn as described above.

## ACKNOWLEDGMENTS

This work was supported with funds from the National Institute of Health/National Institute of Allergy and Infectious Diseases grant R21AI156762 and R21AI166832. The content is solely the responsibility of the authors and does not necessarily represent the official views of the funding agencies. The funders had no role in study design, data collection and analysis, decision to publish, or preparation of the manuscript. “And when you asked for light, I set myself on fire”. - Chris Cornell.

## SUPPLEMENTARY MATERIAL

**FIGURE S1.**
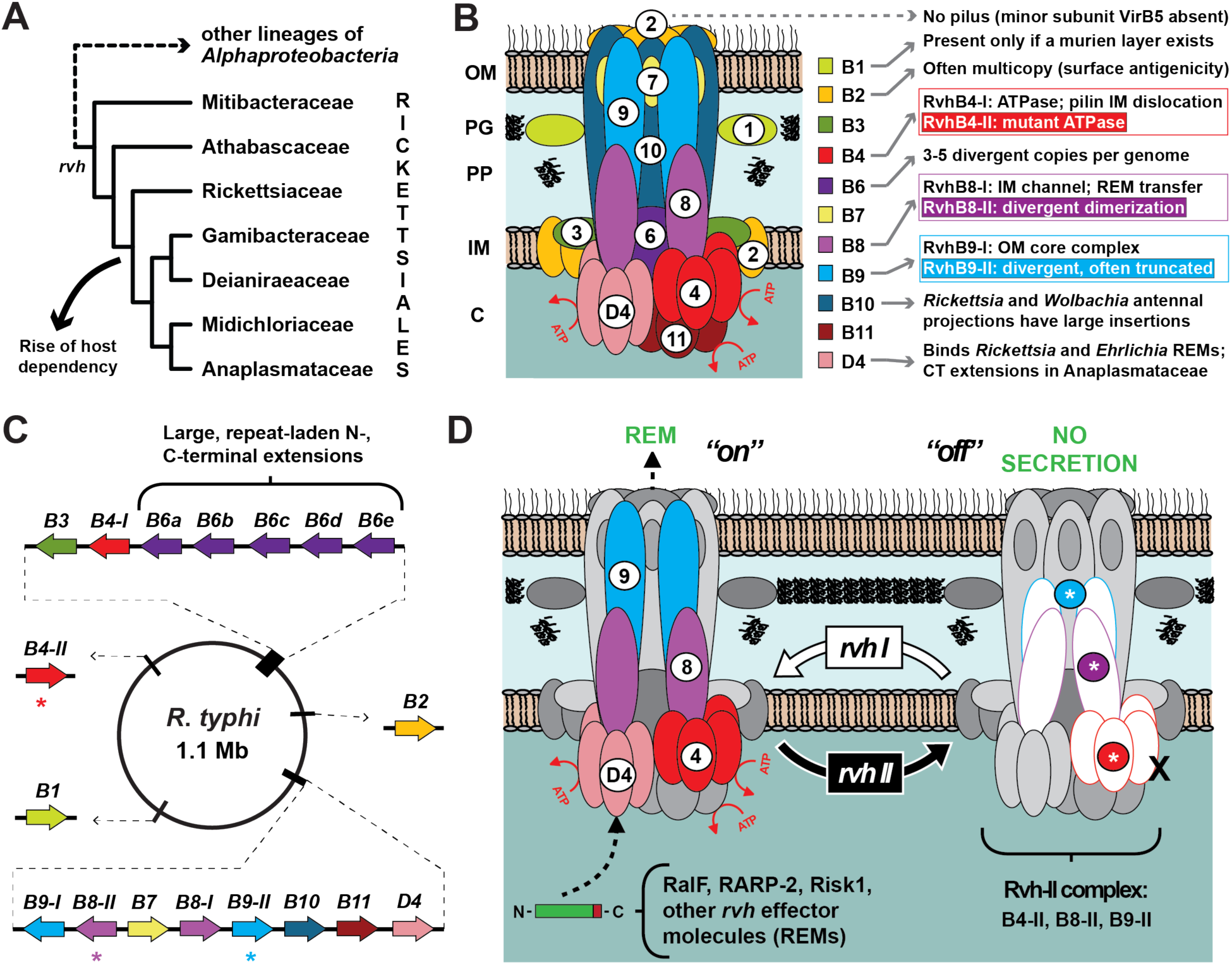
Characteristics of the Rickettsiales *vir* homolog (*rvh*) type IV secretion system (T4SS). **(A)** Schema depicting genome-based Rickettsiales phylogeny estimation, after Schön *et al.* ^28^. Host dependency evolved after the divergence of basal families Mitibacteraceae and Athabascaceae. The *rvh* T4SS was present in the Rickettsiales ancestor. **(B)** Description of the general *rvh* T4SS characteristics, summarized from prior studies ^37, 41, 44, 88^. **(C)** *rvh* genes are arrayed in clustered islets or single transcriptional units throughout *Rickettsia* genomes (e.g., *R*. *typhi* str. Wilmington). Asterisks denote genes for RvhB9-II, RvhB8-II and RvhB4-II. Similar *rvh* gene arrangements are found in other Rickettsiales genomes ^37^. **(D)** Proposed model for conserved *rvh* gene duplication (*rvhB9*, *rvhB8* and *rvhB4*) based on divergent characteristics of RvhB9-II, RvhB8-II and RvhB4-II ^43, 44^.

**FIGURE S2.**
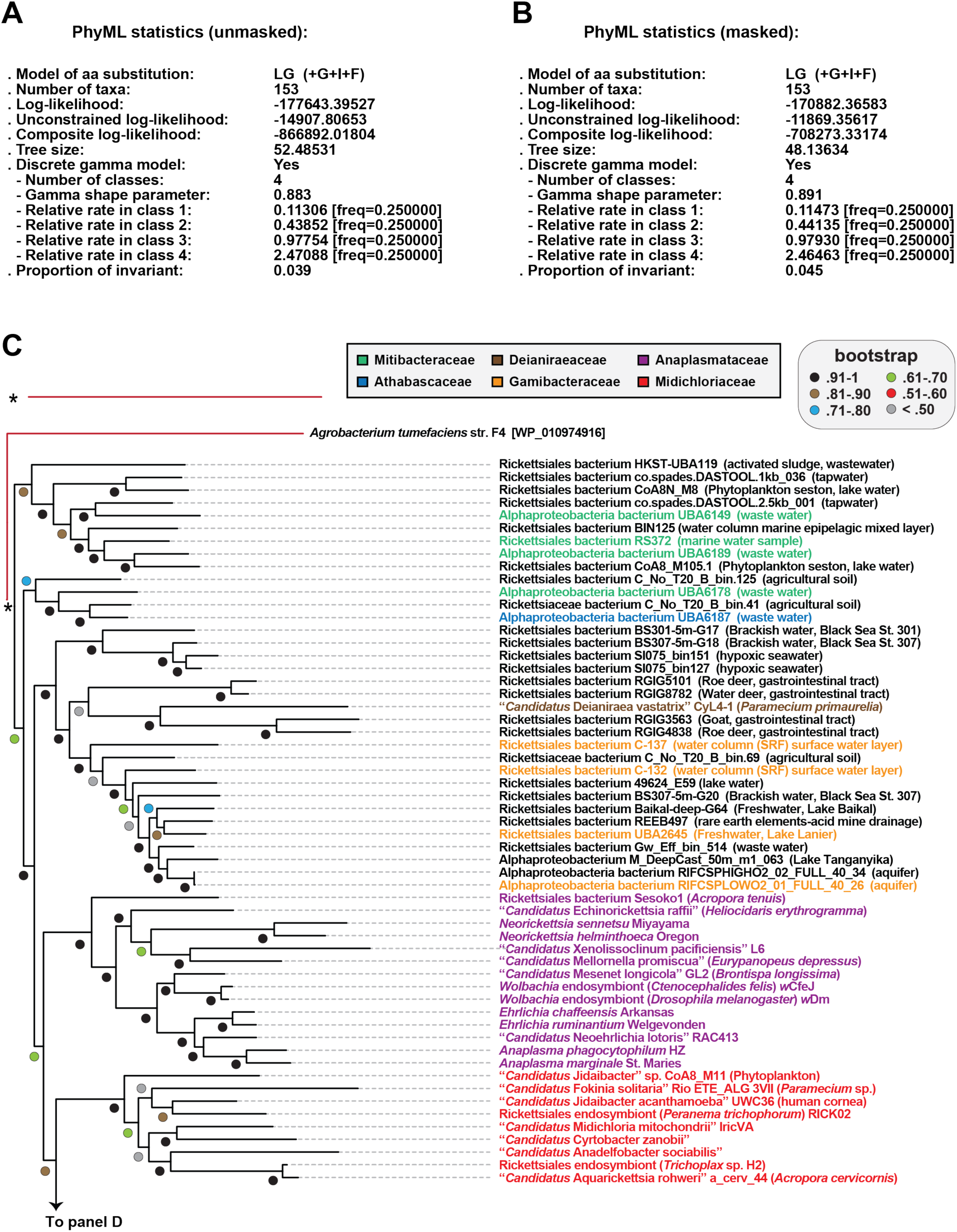
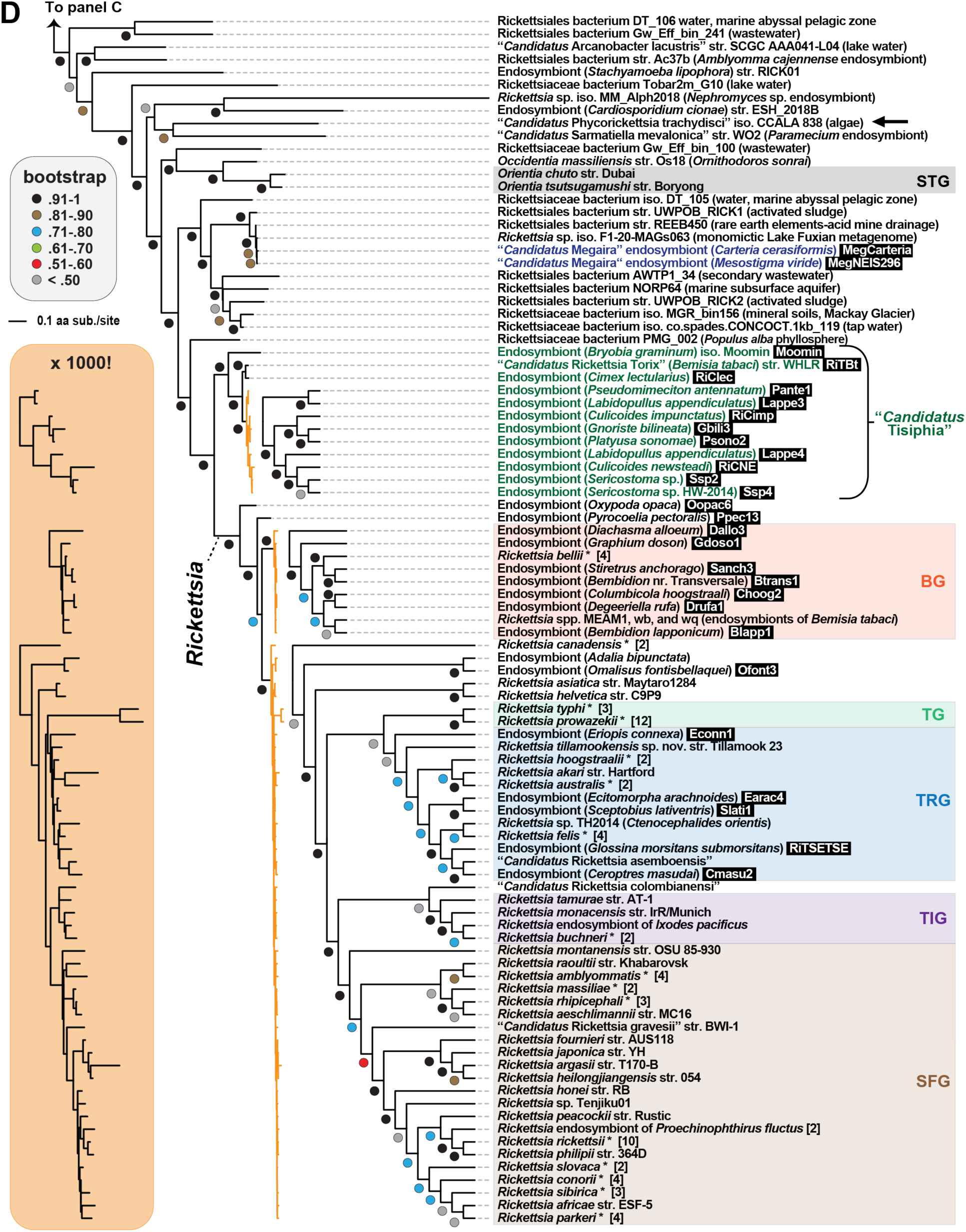
RvhB4-based phylogeny estimation. Phylogeny estimated from concatenated alignments of RvhB4-I and RvhB4-II proteins from 153 rickettsial genome assemblies. *Agrobacterium tumefaciens* str. F4 VirB4 was used as an outgroup. See **Table S1** for sequence information. RvhB4-I and RvhB4-II proteins were separately aligned using MUSCLE (default parameters) with both alignments concatenated (1974 positions). TRIMAL ^181^ was used to create a second dataset with less conserved positions masked (1613 positions). For both unmasked and masked alignments, a maximum likelihood-based phylogeny was estimated with PhyML ^182^, using the Smart Model Selection ^183^ tool to determine the best substitution matrix and model for rates across aa sites (LG (G+I+F) for both alignments). Branch support was assessed with 1,000 pseudo-replications. **(A)** Statistics for the phylogeny estimated on the unmasked alignment, which is shown as a phylogram in panels **C** and **D**, and as a cladogram in **FIG. 1** and **FIG. 2**. **(B)** Statistics for the phylogeny estimated on the masked alignment. This tree is congruent in topology and branch support to the tree generated on the unmasked alignment. **(C)** Non-Rickettsiaceae lineages. **(D)** Rickettsiaceae lineages. “*Candidatus* Sneabacter namystus” (arrow), which lacks *rvh* genes but carries a type VI secretion system T6SS (see FIG. S3) is shown on the phylogram per prior phylogeny estimation ^184, 185^. Black boxes provide short names for 29 MAGs from Davison *et al*. ^72^ (NOTE: the green colored clade comprises genus *Tisiphia* though “*Rickettsia*” reflects NCBI taxonomy as of Feb. 26^th^, 2023). Asterisks, multiple genome assemblies for a species. STG, Scrub Typhus Group; BG, Bellii Group; TRG, Transitional Group; TG, Typhus Group; TIG, Tamurae-Ixodes Group; SFG, Spotted Fever Group. For **C** and **D**, taxa not previously described at the family level are in black text.

**FIGURE S3.**
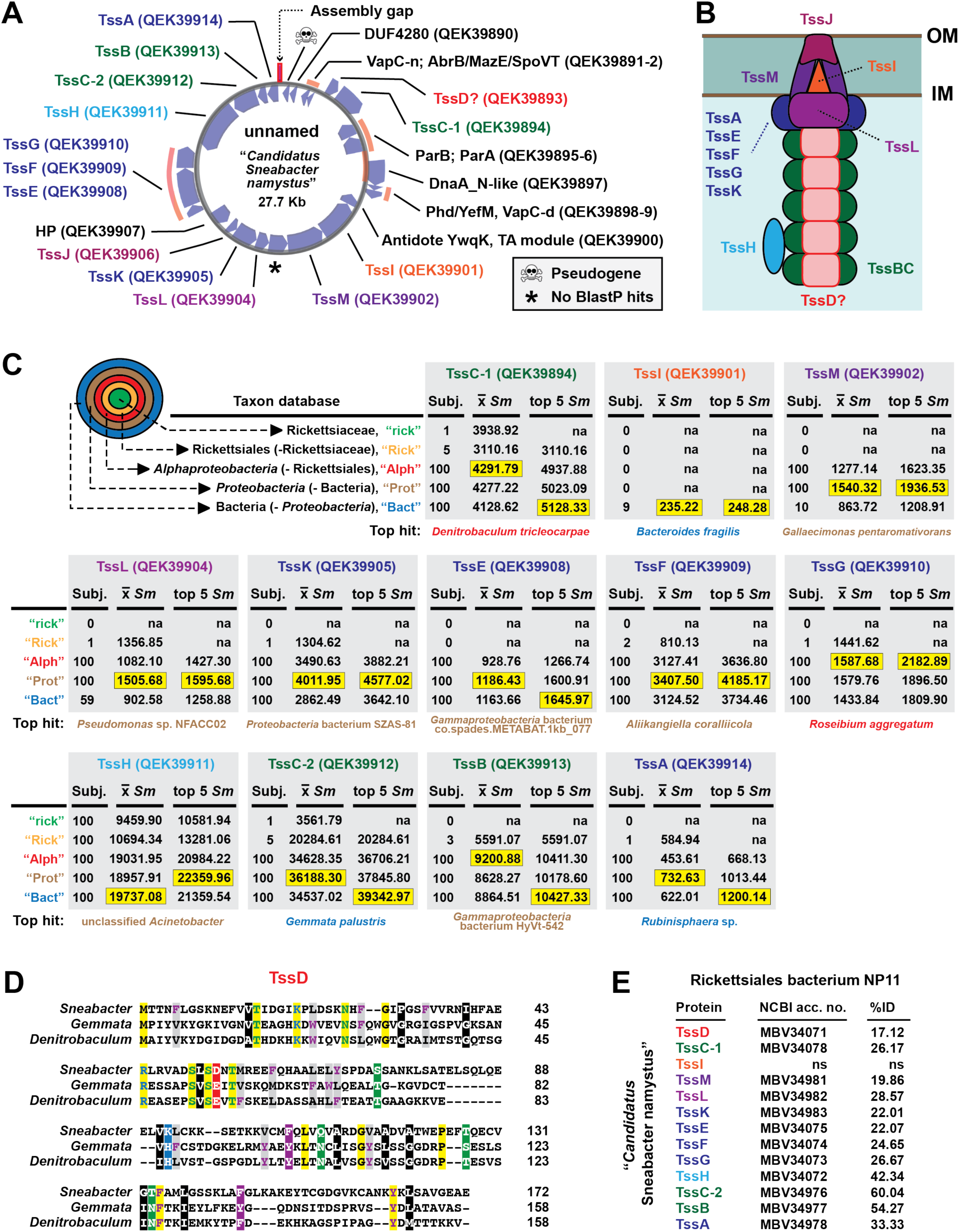
Non-orthologous replacement of secretion machines in Rickettsiales. **(A)** Predicted proteins encoded on the “*Candidatus* Sneabacter namystus” unnamed plasmid, which mostly encodes type VI secretion system (T6SS) components. Protein colors depict their position in panel **B**. Plasmid map created with Proksee (https://proksee.ca/). **(B)** Model of a typical T6SS. The assignment of QEK39893 as a TssD protein is putative (see panel **D**). **(C)** HaloBlast analysis of 12 “*Candidatus* S. namystus” T6SS proteins. BlastP searches were performed against specified NCBI taxon databases (see **TABLE S3** for retrieved sequences). Highlighting: *Sm* score (= *b* * *I* * *Q*, where *b* is the bitscore of the match, *I* is the percent identity, and *Q* is the percent length of the query that aligned) ^26, 73, 202^. **(D)** Alignment of the putative TssD protein of “*Candidatus* S. namystus” with annotated TssD proteins from *Gemmata palustris* (Planctomycetia, WP_210656154) and *Denitrobaculum tricleocarpae* (Rhodospirillales, WP_142896164). Alignment done with MUSCLE ^186^ using default parameters. Amino acid coloring is described in the **FIG. 3** legend. **(E)** Detection of a second complete T6SS in the metagenome assembly Rickettsiales endosymbiont NP11 (NCBI BioSample SAMN07620291) ^203^. Another metagenome assembly, Rickettsiales endosymbiont EAC13 (SAMN07620031) ^203^ contained fewer T6SS genes. Neither assembly harbors *rvh* genes (data not shown), indicating that other Rickettsiales species may utilize a T6SS in place of a T4SS.

**FIGURE S4.**
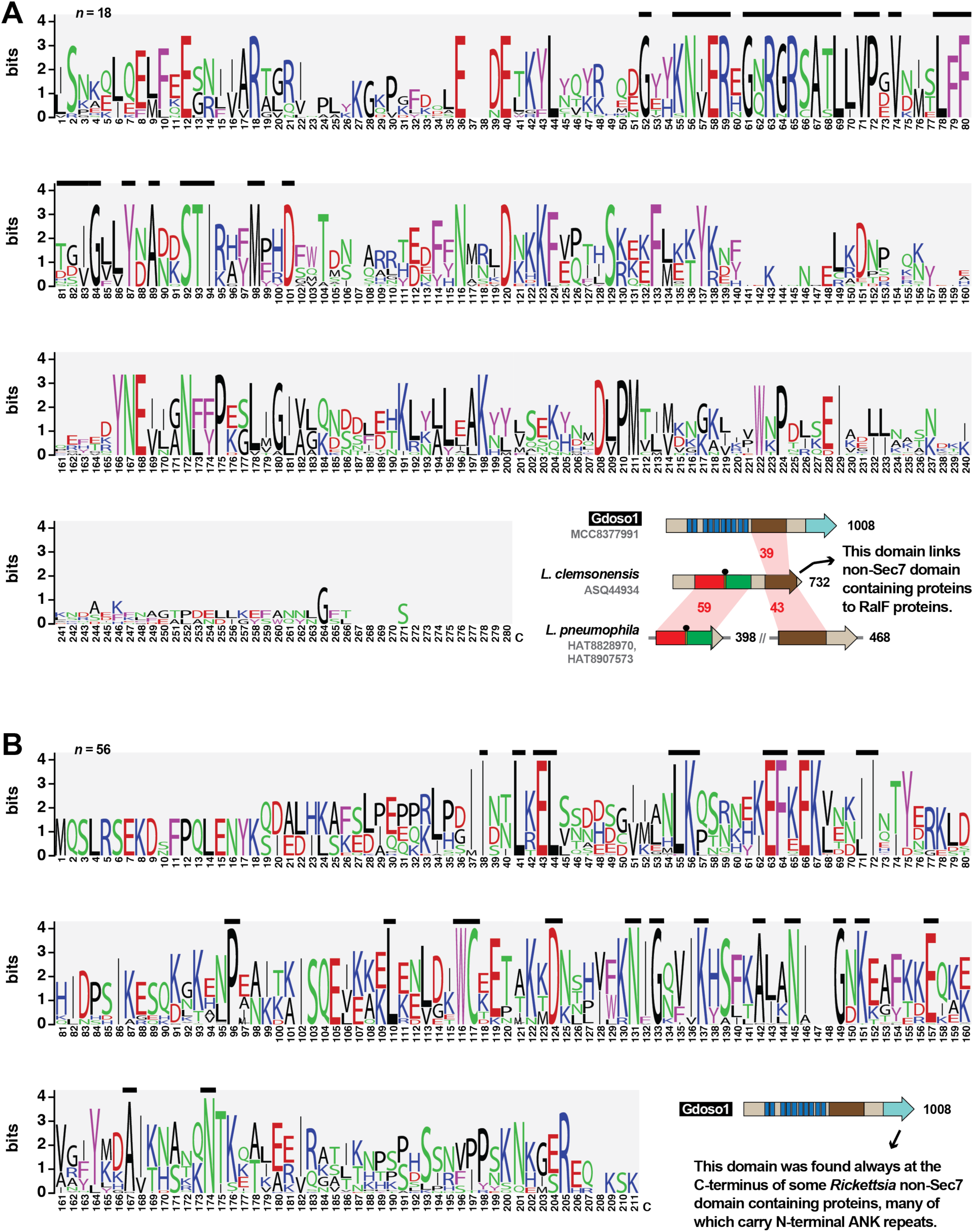
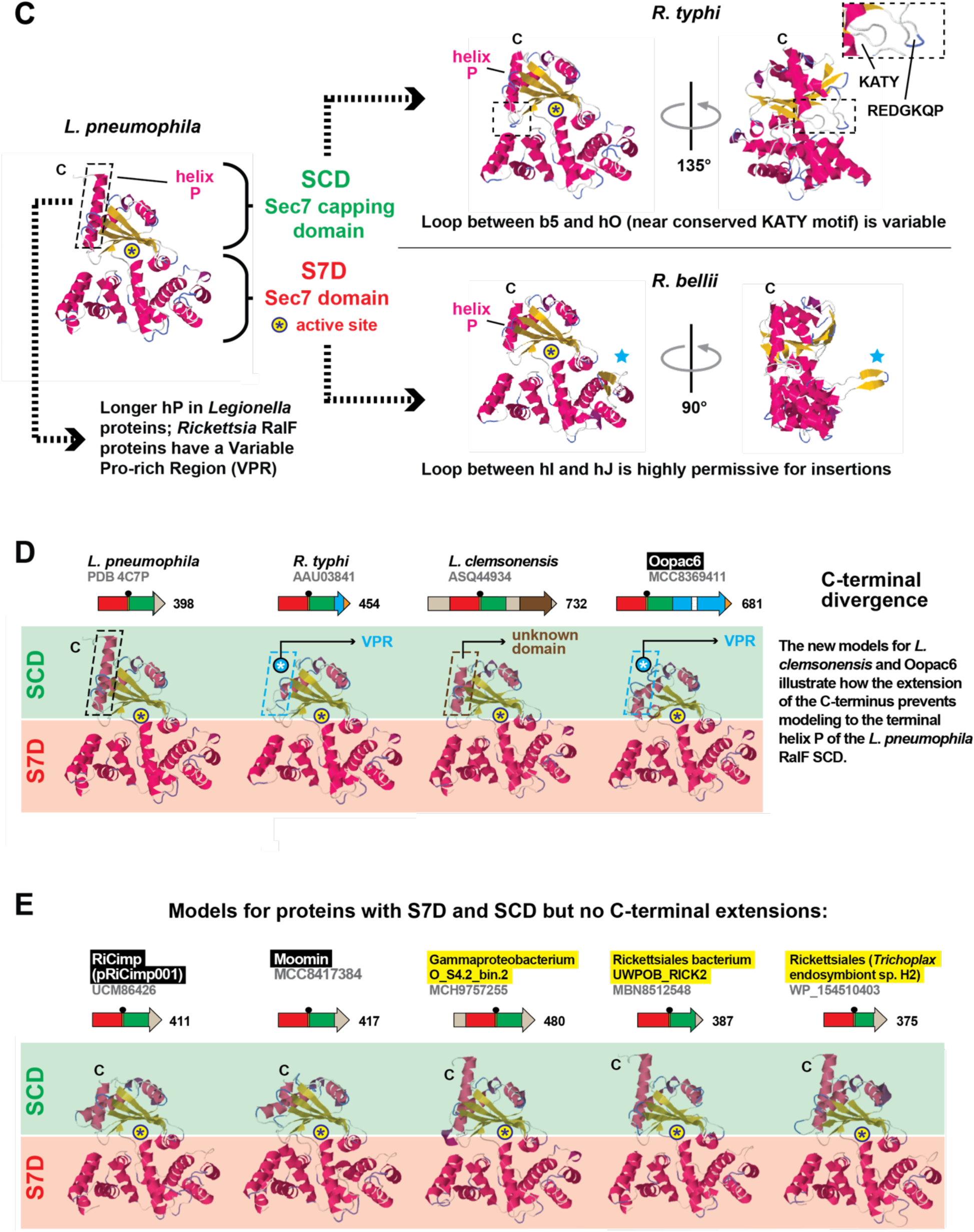
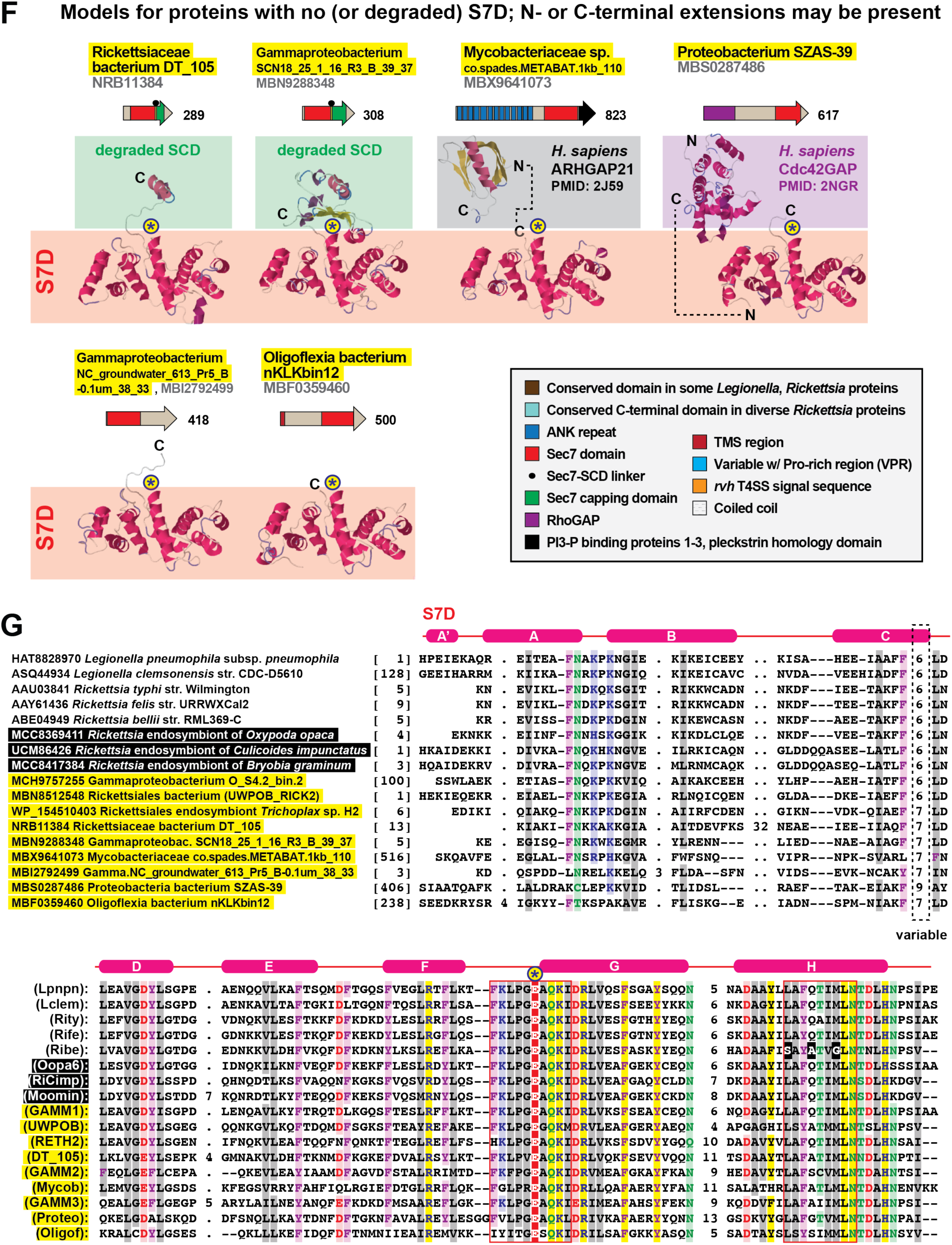
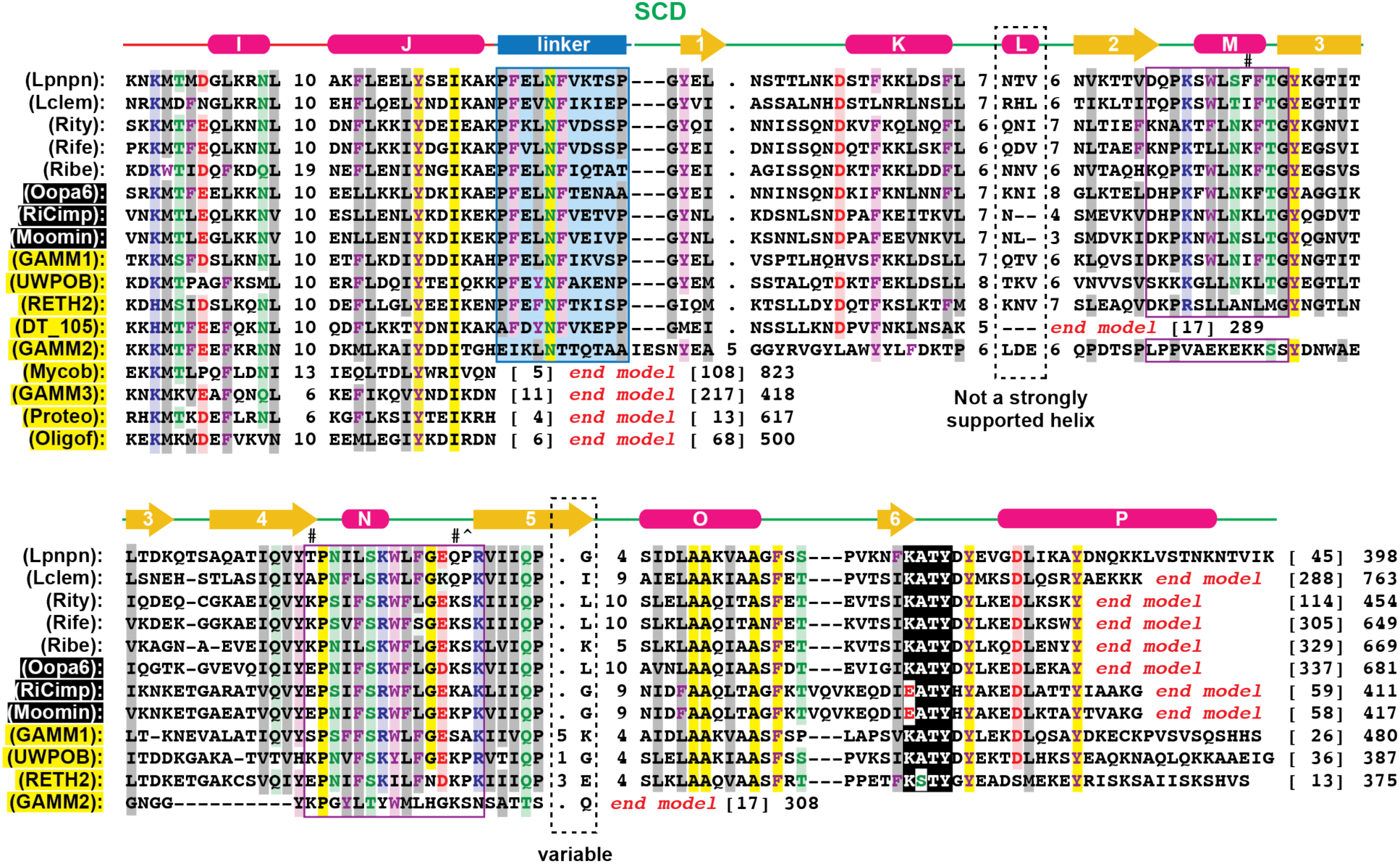
Phylogenomics and bioinformatics analyses of RalF proteins. Amino acid coloring is described in the **FIG. 3** legend. Black boxes provide short names for MAGs from Davison *et al*. ^72^. These and additional newly discovered RalF-like proteins (highlighted yellow) substantially expand the prior RalF diversity. **(A, B)** Sequence logos ^187^ illustrating conservation within two novel domains present in some *Legionella* and *Rickettsia* proteins. Sequence information is provided in **Table S2**. Sequences were aligned with MUSCLE ^186^ using default parameters. These two domains are carried in a single protein (MCC8377991) from *Rickettsia* endosymbiont of *Graphium doson* (Gdoso1). **(A)** Sequence conservation (*n* = 18) within the Gdoso1 MCC8377991 central domain. **(B)** Sequence conservation (*n* = 56) within the Gdoso1 MCC8377991 C-terminal domain. **(C)** Structural characteristics of the RalF S7D-SCD architecture. ***Left*:** *L*. *pneumophila* RalF structure (PDB 4C7P) ^85^. The delineation of the Sec7 domain (S7D, red) and Sec7-capping domain (SCD, green) is shown, with an approximation of the active site Glu (asterisk). ***Right*:** regions of variability within the predicted structures of *R*. *typhi* str. Wilmington (RT0362) and *R*. *bellii* str. RML369-C (RBE_0868) RalF proteins. Modeling done with Phyre2 ^192^. *Rickettsia* full length RalF proteins contain an extended C-terminal domain relative to *L*. *pneumophila* RalF ^55^. **(D-F)** Gallery of predicted structures for diverse RalF and RalF-like proteins. All protein domains are described in the gray inset. **(G)** Sequence comparison of diverse RalF and RalF-like proteins (S7D and SCD only). An initial alignment with MUSCLE (default parameters) was manually adjusted with reference to the Phyre2 structure models to *L*. *pneumophila* RalF (PDB 4C7P). The secondary structure of *L*. *pneumophila* RalF is superimposed over the alignment. Conserved residues are highlighted yellow. S7D: two highly conserved regions (motif 1 and motif 2) that together form the Sec7 active site are boxed in red ^204^; the active site Glu of Motif 1, which is essential for Arf1 recruitment to the LCV ^84^ and Arf6 recruitment to the plasma membrane during *Rickettsia* invasion ^147^, is noted with an asterisk; divergent residues within motif 2 of *R*. *bellii* RalF are colored black. SCD: aromatic clusters comprising the membrane sensor region are enclosed in purple boxes; following previous mutagenesis analysis of the SCD ^86^, # denotes residues in *L*. *pneumophila* RalF permuted to the corresponding residues in *R*. *prowazekii* RalF (and vice versa) and ^ denotes residues in *L*. *pneumophila* RalF permuted to the corresponding residues in *R*. *prowazekii* RalF but not reciprocated. The conserved KATY motif, which contacts the S7D remote from the active site and is thought to function as a hinge for the conformational change that activates RalF ^86^, is colored black.

**FIGURE S5.**
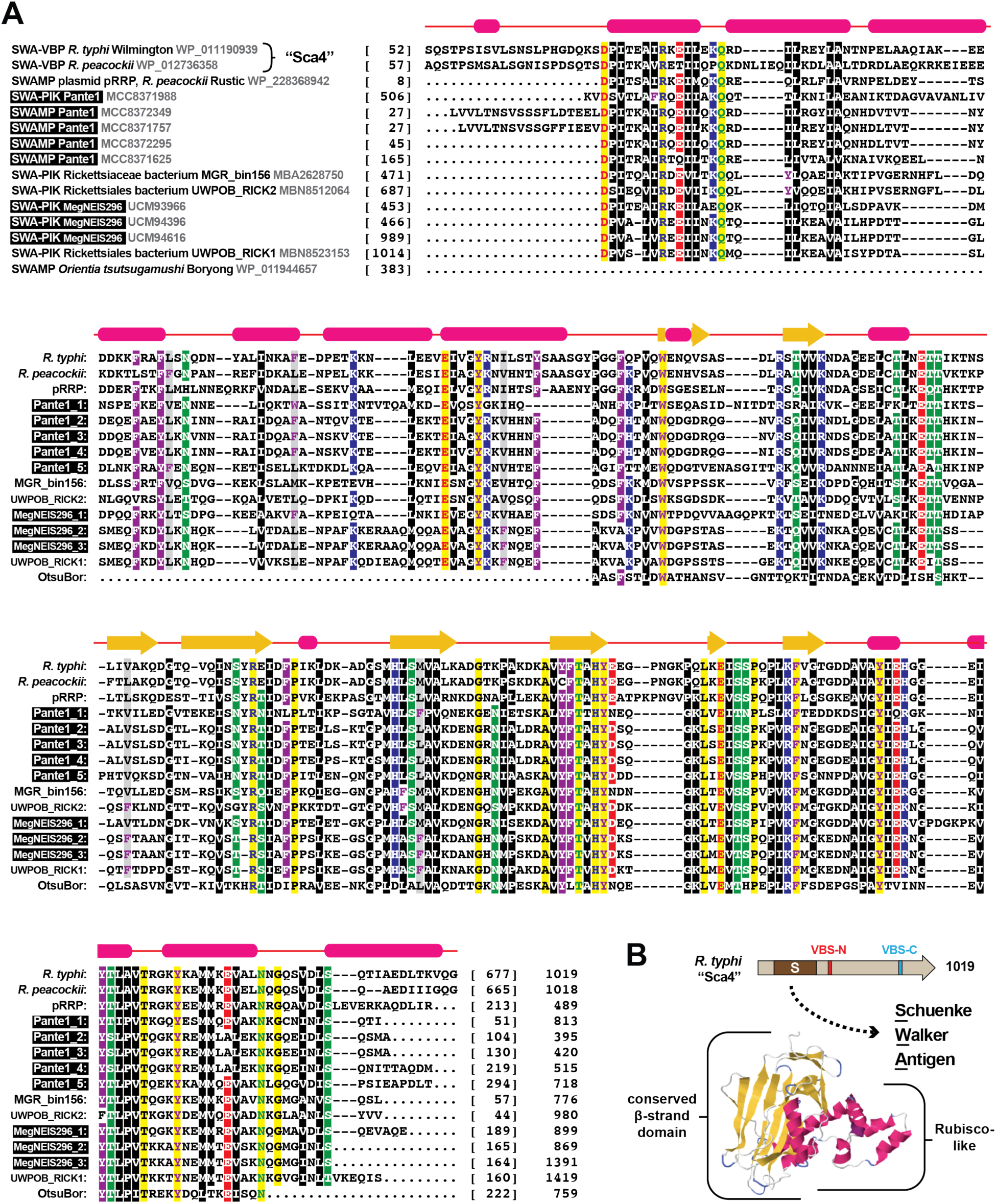
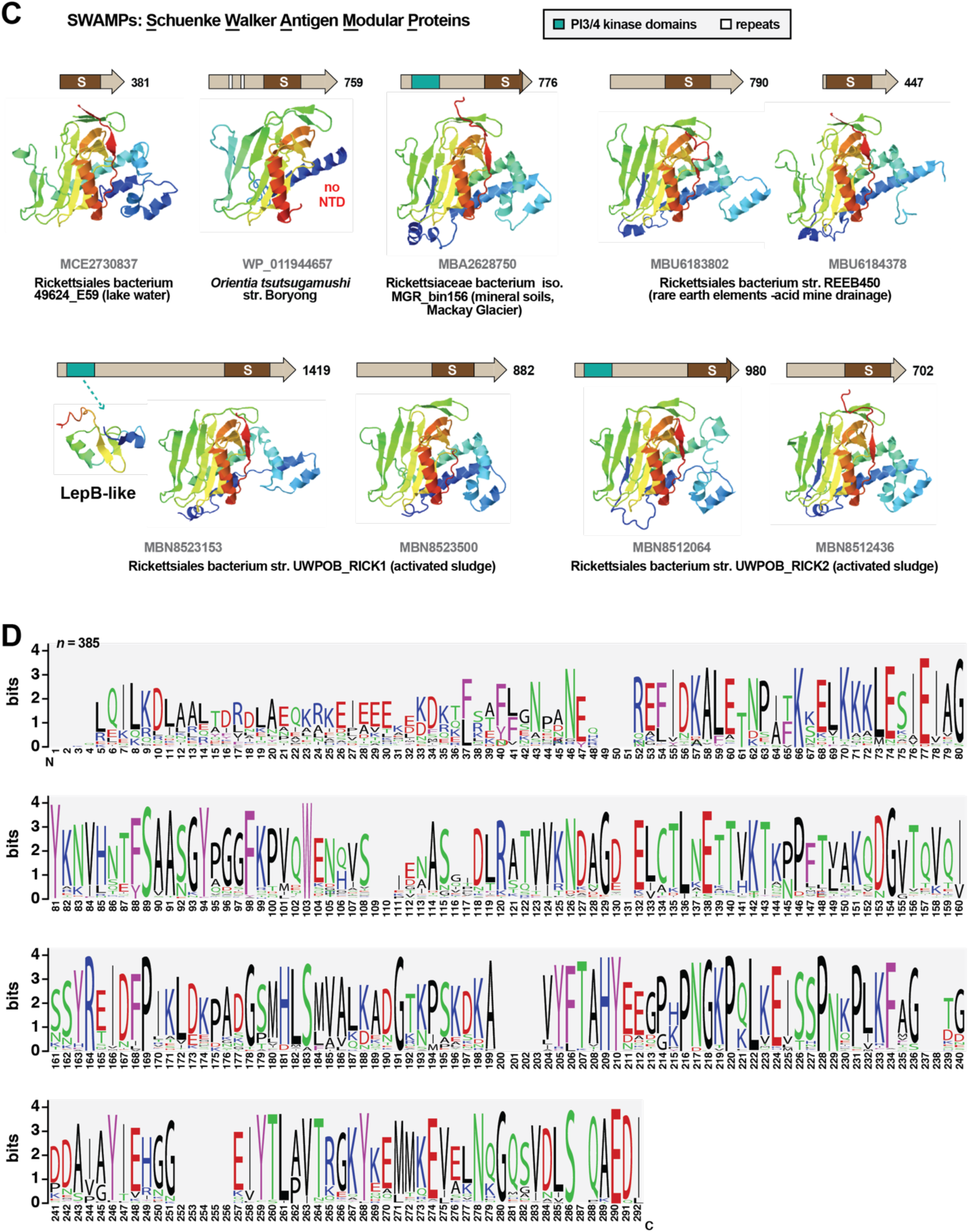
The N-terminal domain of *Rickettsia* Sca4 proteins is recurrent and widespread in other rickettsial proteins. Amino acid coloring is described in the **FIG. 3** legend. Black boxes provide short names for MAGs from Davison *et al*. ^72^. **(A)** Alignment of the Schuenke Walker Antigen (SWA) domains of select *Rickettsia* Sca4 proteins with analogous domains in SWA-Risk2 chimeras (SWA-PIK) and SWA modular proteins (SWAMPs). Domains were retrieved from BlastP searches against the NCBI nr protein database using the *R. typhi* SWA domain as a query. Sequences were aligned with MUSCLE ^186^ using default parameters. Structural assignment above alignment corresponds to the modeling of *Rickettsia* Sca4 N-terminal domains to the *R*. *rickettsii* SWA domain (PDB ID: 4LQ8) ^111^. **(B)** Illustration of the SWA domain of *R. typhi* Sca4 and the two Vinculin Binding Sites (VBS); VBS-C of *R*. *rickettsii* interacts with the head domain of human vinculin ^776^. **(C)** Gallery of diverse architectures for select SWAMPs and predicted structures of SWA domains. Phyre2 ^192^ was used to model all SWA structures to the *R*. *rickettsii* SWA domain, as well as the PIK domain of Rickettsiales bacterium str. UWPOB_RICK1 to a portion of the LepB protein of *L. pneumophila* (PDB ID: 4JW1) ^199^. Domains on schemas were predicted with SMART ^188^. **(D)** Sequence logos ^187^ depict a consensus from an alignment of all 385 SWA domains retrieved from a BlastP searches against the NCBI nr protein database using the *R. typhi* SWA domain as a query (sequence information in **Table S2**).

**FIGURE S6.**
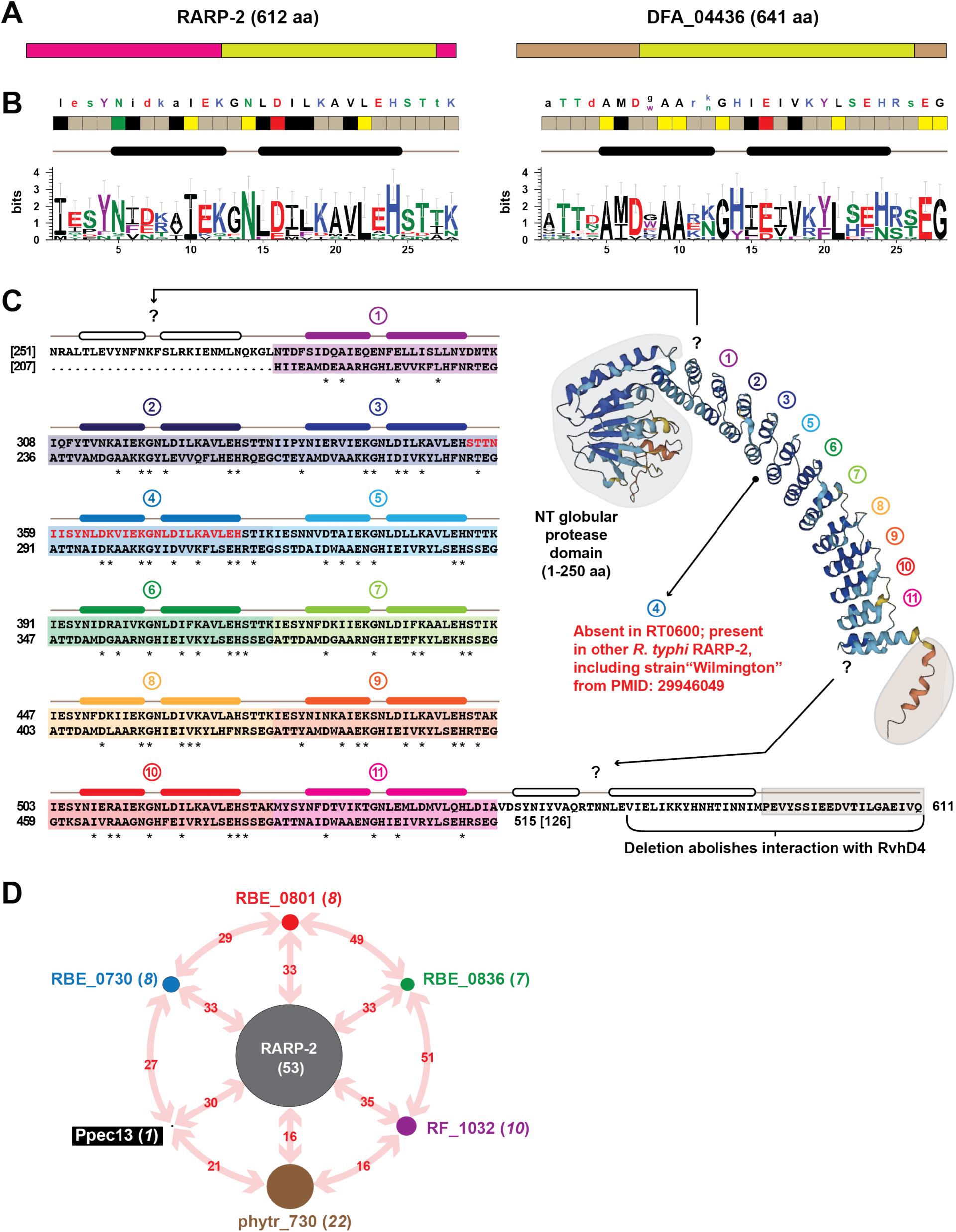
Characteristics of *Rickettsia* Ankyrin Repeat 2 (RARP-2) proteins. **(A)** Comparison of the domain architecture for *Rickettsia* RARP-2 proteins (*R*. *typhi* str. Wilmington as an exemplar, NCBI acc. no. AAU04065) and the ANK repeat-containing protein DFA_04436 (XP_004360169) of the cellular slime mold *Cavenderia fasciculata* (Eumycetozoa; Dictyostelia; Acytosteliales). **(B)** Sequence logos ^187^ constructed to illustrate the conservation between ANK repeats within *R*. *typhi* RARP-2 and D DFA_04436 and between the two models. Repeats were manually stacked and visualized with AliView 1.28.1 ^205^. Amino acid coloring is described in the **FIG. 3** legend. **(C)** Structure of the *R*. *typhi* RARP-2 C-terminal ANK repeat domain. **(*left*)** Alignment of the entire ANK repeat-containing domains between *R*. *typhi* RARP-2 and *C. fasciculata* DFA_04436. Alignment constructed using MUSCLE ^186^ (default parameters). Helices above ANK repeats are derived from the conserved ANK repeat structure ^117^. Red sequence illustrates the fourth ANK repeat present in RARP-2 of *R*. *typhi* str. TH1527 (AFE54444), *R*. *typhi* str. B9991CWPP (AFE55282) and the laboratory strain named “Wilmington” that differs from AAU04065 and remains unpublished ^57^. **(*right*)** Prediction of entire RARP-2 structure using Alphafold ^194, 195^. Additional ANK-like folds flanking the 11 ANK repeats predicted by comparative analyses are noted with question marks. **(D)** Summary of divergent RARP-2 (dRARP-2) proteins. N-terminal protease domains from each of the six sequences in the outer circle were used as a query in BlastP searches and determined to have greater similarity to a cohort (number in parentheses and size of dot) or itself only (Pyropec, or *Rickettsia* endosymbiont of *Pyrocoelia pectoralis*, a MAG from Davison *et al*. ^72^) versus other query sequences or any RARP-2 protein (center). All sequence information is provided in **Table S2**.

**FIGURE S7.**
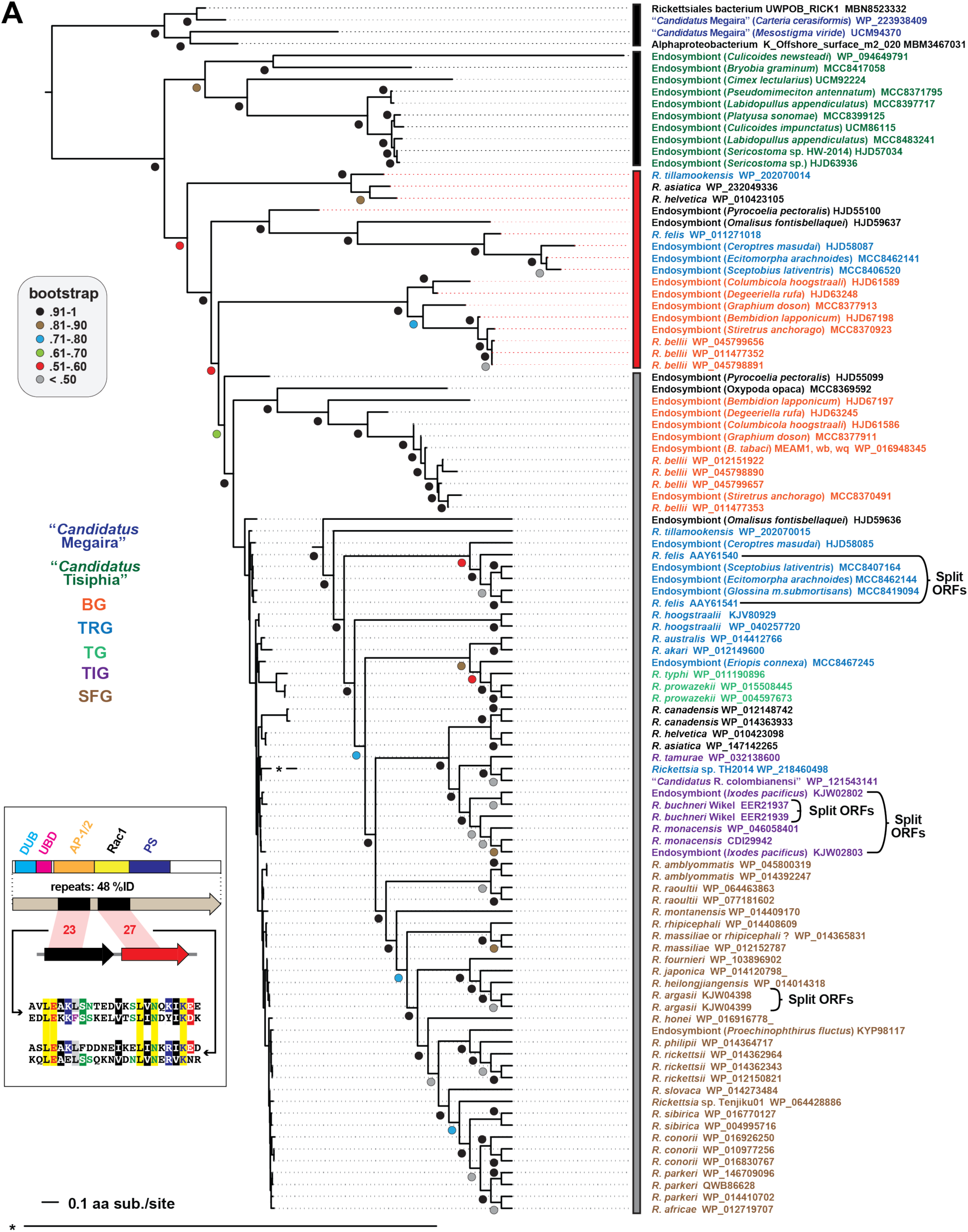
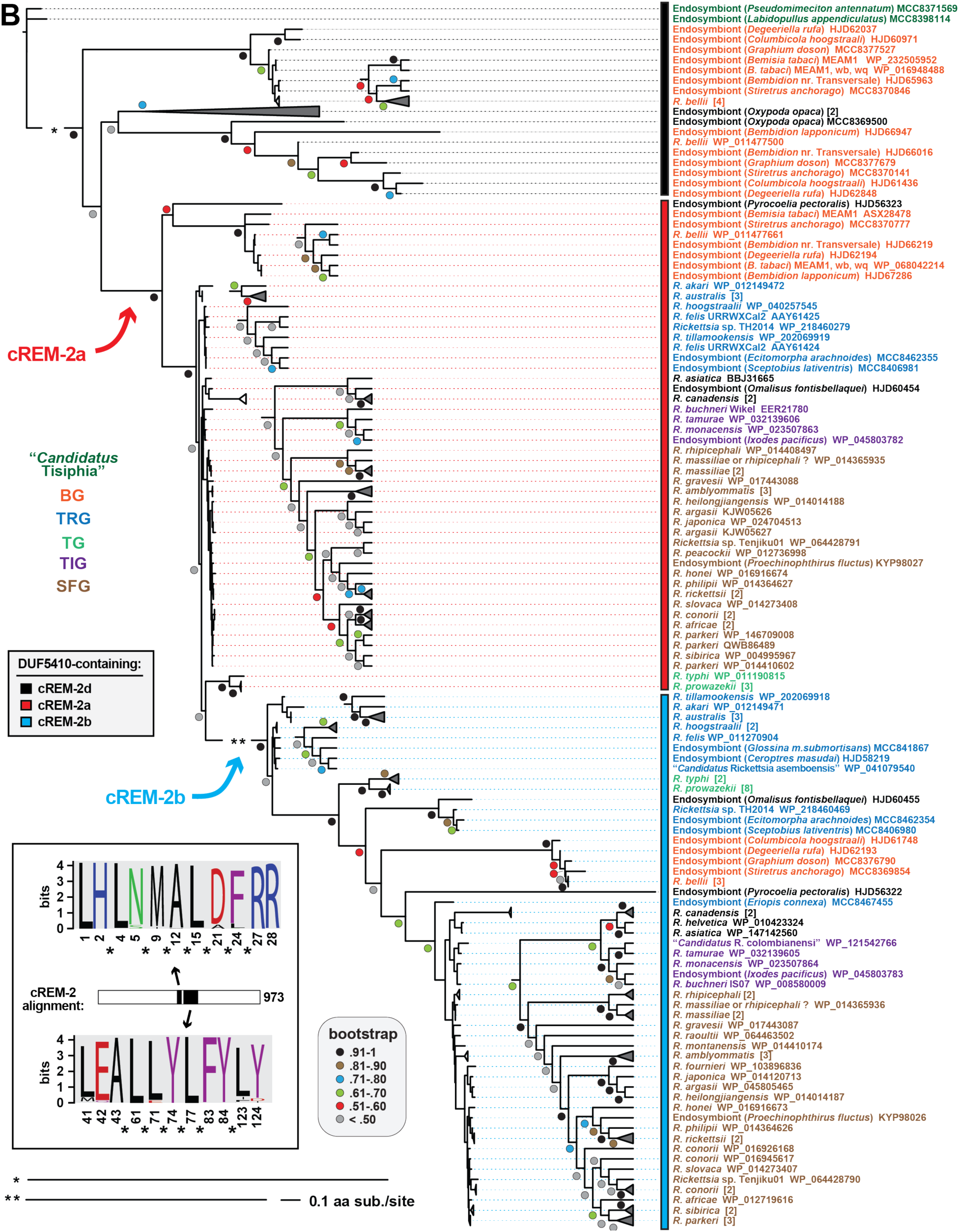
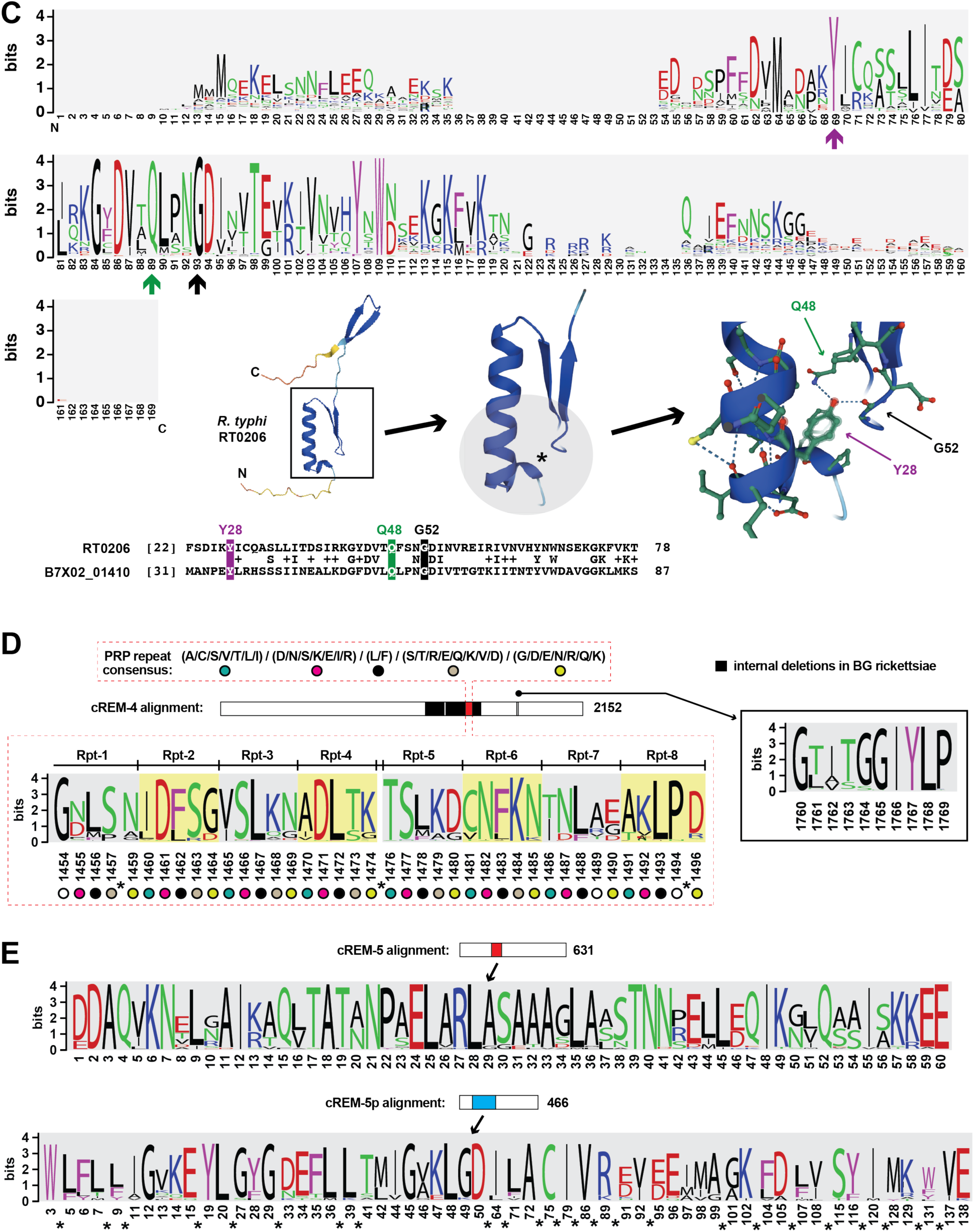
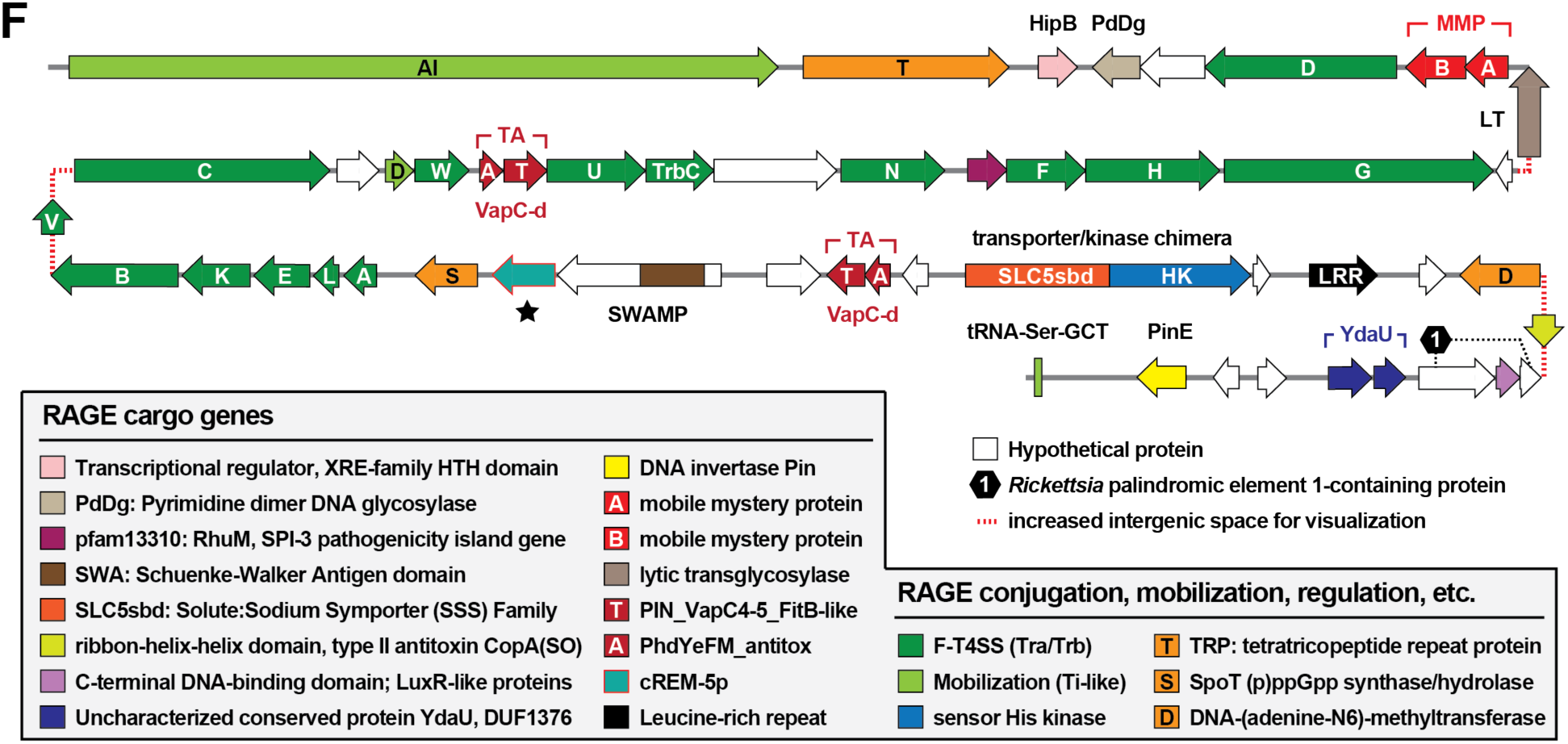
Phylogenomics analyses of candidate REMs (cREM-1-cREM-5). Protein information is provided in **Table S2**. All alignments done with MUSCLE (default parameters) ^186^ with conservation analyzed using WebLogo ^187^. Amino acid coloring is described in the **FIG. 3** legend. Black boxes provide short names for MAGs from Davison *et al*. ^72^. **(A)** cREM-1 proteins are minimized in architecture relative to ancestral forms. Proteins with high similarity to RT0435 (cREM-1) are mostly conserved in *Rickettsia* genomes yet highly diverse (gray), sometimes duplicated and tandemly arrayed (red) and often components of larger modular proteins (cREM-1d). Inset illustrates cREM-1 similarity to tandem repeats in the scrub typhus effector OtDUB (CAM80065), which carries multiple eukaryotic-like domains ^5, 123, 201^ (described in **FIG. 7**). For brevity, alignment of a small conserved region shared across OtDUB repeat 1 (383-406), OtDUB repeat 2 (645-668), Blapp1 HJD67197, and Blapp1 HJD67198 is shown. Phylogeny estimation of 102 cREM-1 proteins indicates diversification of larger CREM-1 domain modular proteins and streamlining to a single cREM-1 protein in most *Rickettsia* genomes. Alignment was not masked (1544 total sites, 38.34% invariant). A maximum likelihood-based phylogeny was estimated with PhyML ^182^, using the Smart Model Selection ^183^ tool to determine the best substitution matrix and model for rates across aa sites (VT +G+F). Branch support was assessed with 1,000 pseudo-replications. Log likelihood of tree: −34728.6. **(B)** cREM-2 proteins diversified from an ancient gene duplication. Proteins with high similarity to RT0352 (cREM-a, red) and RT0351 (cREM-2b, light blue) are tandemly arrayed and mostly conserved in *Rickettsia* genomes, yet highly diverse from ancient forms (cREM-2d). Inset illustrates the conserved central region of cREM-2 proteins. Estimated phylogeny indicates cREM-2b proteins are more divergent from cREM-2d proteins. Alignment was not masked (973 total sites, 43.4% invariant). Phylogeny estimated as described in panel **A** (final model LG +G). Branch support was assessed with 1,000 pseudo-replications. Log likelihood of tree: −25322.17. **(C)** cREM-3 proteins are highly conserved and present in other proteobacterial assemblies. These proteins are typically annotated as Pfam PF10877 (DUF2671), which states they are restricted to *Rickettsia* spp. This sequence logo is for 107 non-redundant proteins obtained from searches against ‘Rickettsiales’, with proteins aligned using MUSCLE (default parameters). A more complete list of proteins is found in **Table S2**, though more sequences are likely retrievable using HMMER searches. At bottom, a pairwise alignment between *R*. *typhi* RT0206 and the most divergent subject retrieved from a BlastP search excluding ‘Rickettsiales’ is shown (hypothetical protein B7X02_01410 from Rhodospirillales bacterium 12-54-5, OYW13786.1). These proteins are 30% identical over the match shown. The residues noted with arrows in the sequence logo are shown over the pairwise alignment and below with a structure for *R*. *typhi* RT0206 predicted with Alphafold ^194, 195^. **(D)** cREM-4 proteins harbor a conserved pentapeptide repeat (PR). Schema shows alignment of 82 non-redundant cREM-4 proteins with illustration of the PR consensus sequence at top^206^. A small conserved motif (inset) was also identified. **(D)** cREM-5 and cREM-5p have conserved central regions that lack similarity to proteins outside of *Rickettsia* and *Tisiphia* genomes. **(F)** One copy of cREM-5p from the RiClec (Endosymbiont of *Cimex lectularius*) genome is found on a RAGE transposon. General schema and annotation of RAGE genes follows previous reports ^90, 91, 132^.

**FIGURE S8.**
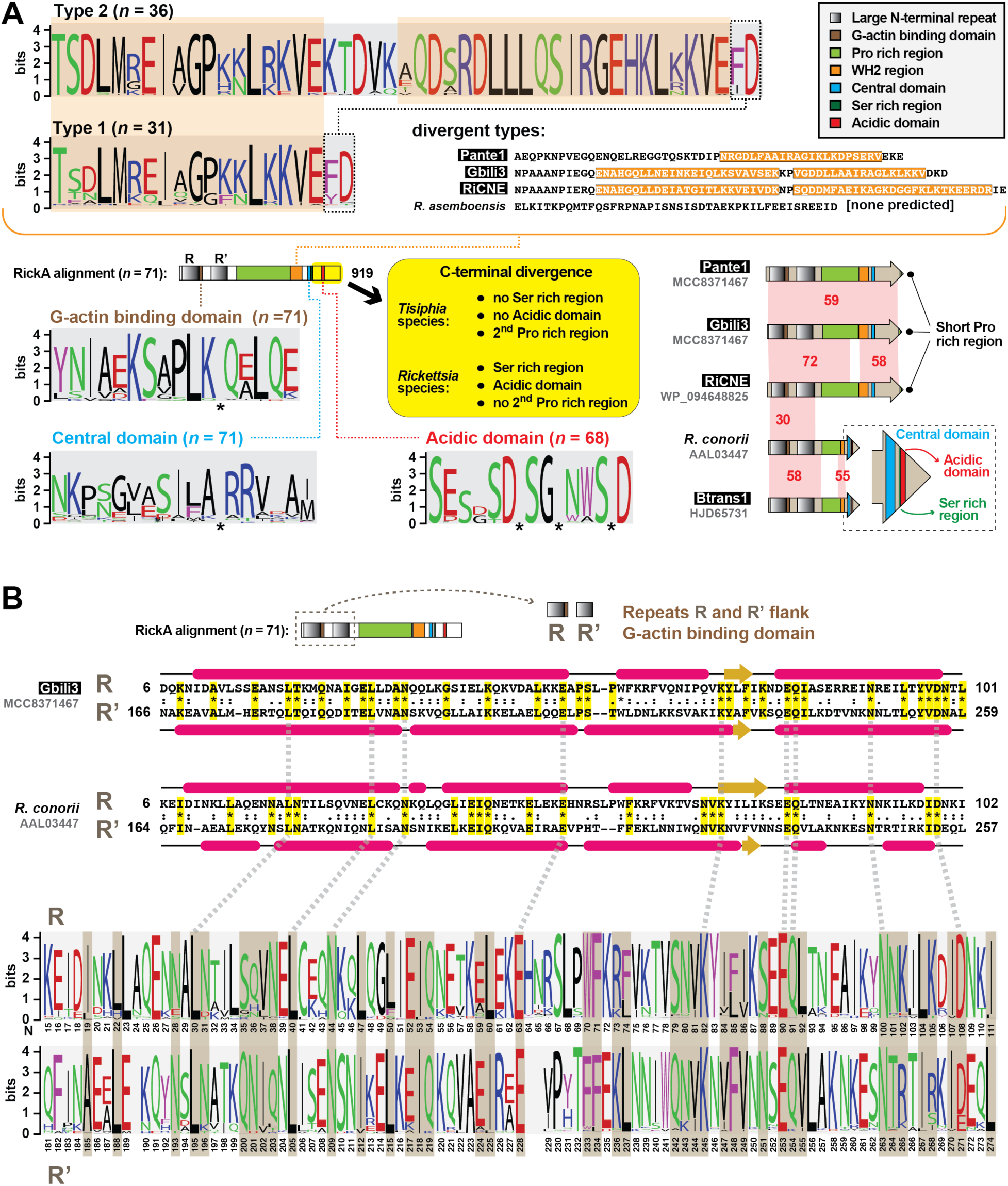
Discovery of novel RickA architecture. Black boxes provide short names for 29 MAGs from Davison *et al*. ^72^. Amino acid coloring is described in the **FIG. 3** legend. Sequence logos constructed with WebLogo 3 ^187^. Sequence information in **Table S2**. **(A)** General architecture of RickA proteins deduced from an alignment of 71 non-redundant sequences using MUSCLE ^186^ (default parameters). Consensus sequences for the WH2 (divided into two types based on one or two motifs) are shown at top, with divergent motifs illustrated. The remaining graphics illustrate the differences between *Tisiphia* and *Rickettsia* RickA C-terminal regions (summarized in the yellow ellipse). **(B)** Identification of a novel N-terminal repeat sequence in all RickA proteins. The repeat was identified by assessing within-protein BlastP matches, which revealed ∼34 %ID between ∼95 aa regions flanking the G-actin-binding domain (depicted in schema at top). Two examples are shown for a *Tishiphia* and *Rickettsia* repeat alignment, with secondary structure predictions indicating high conservation between repeats ^196^. A comparison of the repeats R and R’ all 71 non-redundant sequences is shown at bottom, with residues showing conservation in amino acid type (hydrophobicity, charge, aromaticity) shaded tan. This collectively illustrates that each RickA protein has greater within-repeat similarity than across-protein repeat similarity yet is constrained in overall amino acid type.

**FIGURE S9.**
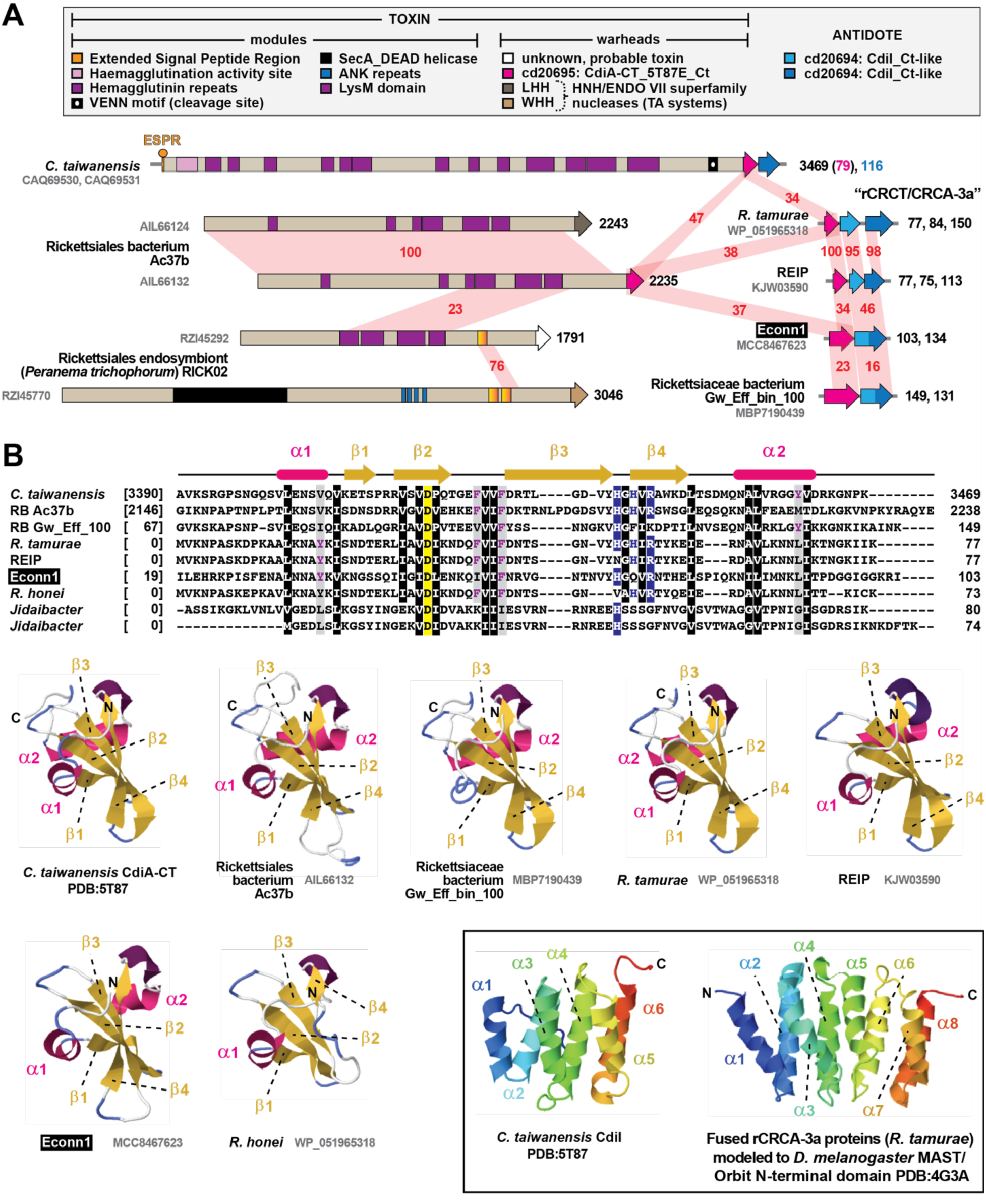
CDI-like/Rhs-like C-terminal toxin/antidote (CRCT/CRCA) modules occur in diverse rickettsial genomes. Black boxes provide short names for 29 MAGs from Davison *et al*. ^72^. Sequence information in **Table S2**. **(A)** Comparison of the *Cupriavidus taiwanensis* str. DSM 17343 CdiA/I module to several rickettsial large modular toxins, as well as select rCRCT/CRCA-3a modules. Domains were predicted with SMART ^188^. Amino acid similarity (%ID, red shading) was assessed using Blastp. **(B)** Structural analysis of a rCRCT/CRCA-3a module. (***top***) Alignment of *C*. *taiwanensis* CdiA with eight rickettsial rCRCT-3a toxins. Amino acid coloring is described in the **FIG. 3** legend. Structural information from *C*. *taiwanensis* CdiA (PDB:5T87) ^207^ is provided at top. Alignment performed using MUSCLE with default settings ^186^. (***bottom***) Modeling with Phyre2 ^192^ of six rCRCT-3a toxins to the CdiA-CT toxin structure of *C*. *taiwanensis* CdiA (PDB:5T87). All threading was done with >95% confidence. The *Jidaibacter* proteins were too divergent to model. Inset: while the *C*. *taiwanensis* module was solved as a co-complex ^207^, Phyre2 modeling could not thread rickettsial rCRCA-3a to the antidote CdiI within the TA co-complex. The best model (46.3% confidence, 12% ID) for *R*. *tamurae* rCRCA-3a was to the structure of *Drosophila melanogaster* MAST/Orbit N-terminal domain PDB:4G3A ^208^, indicating a similar helical bundle and similar topology as *C*. *taiwanensis* CdiI.

**FIGURE S10.**
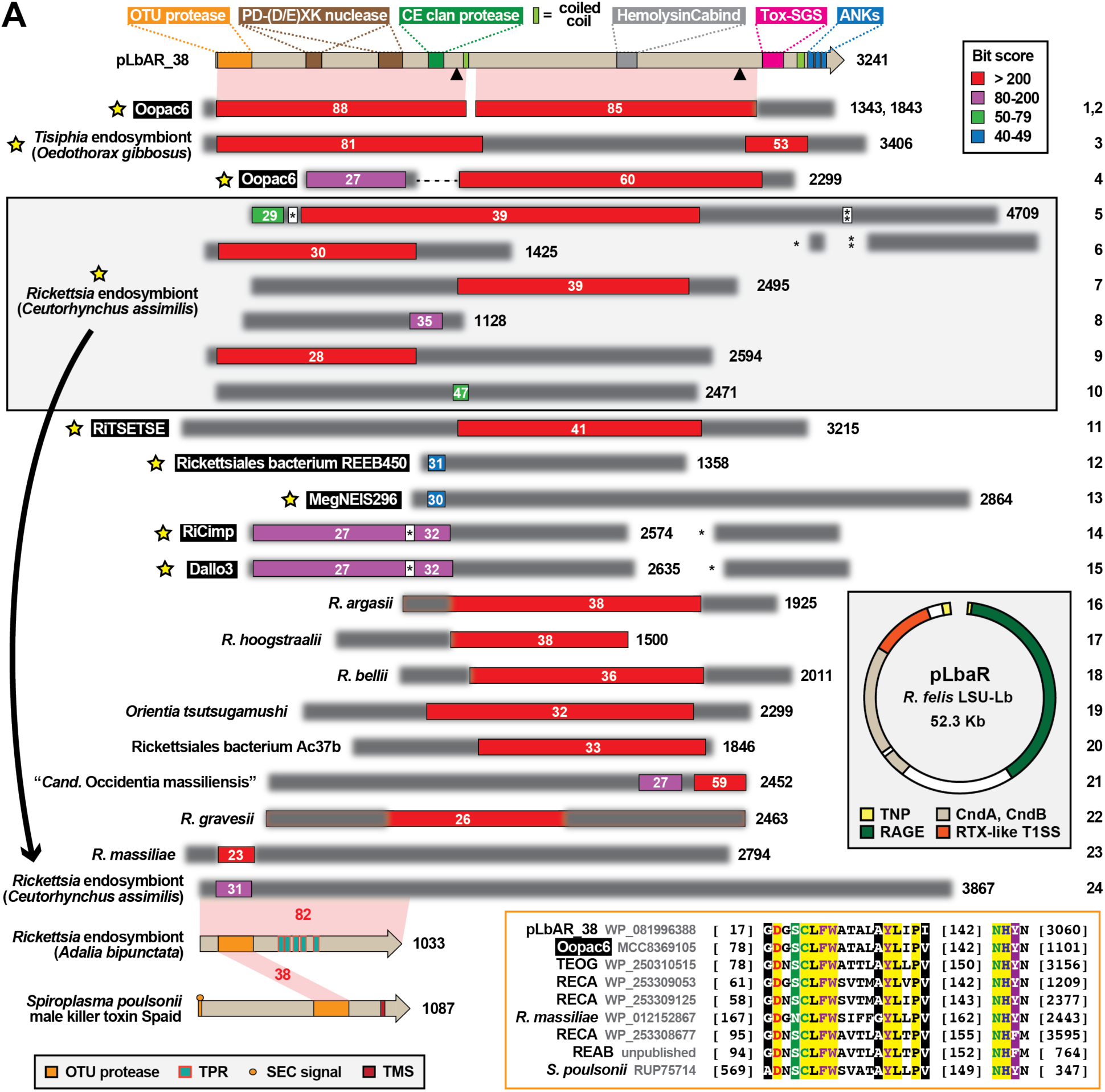
Identification of MAG proteins that have characteristics of reproductive toxins. Black boxes provide short names for 29 MAGs from Davison *et al*. ^72^. Sequence information in **Table S2**. [NOTE: The *Rickettsia* endosymbiont of *Oedothorax gibbosus* (NZ_OW370493) and *Rickettsia* endosymbiont of *Ceutorhynchus assimilis* (NZ_OU906081) assemblies were not included in our other analyses, as manuscripts supporting these assemblies were not published. We included them in this analysis due to their relevance to reproductive parasitism (RP) in *Rickettsia* species]. ***Top*,** architecture of the modular protein pLbAR_38, which is carried on plasmid pLbAR of *Rickettsia felis* str. LSU-Lb (gray inset) ^91^. Black triangles, proprotein convertase cleavage sites ^209^. We refer to this protein and its adjacent predicted antidote (not shown) as a CndA/B module ^173^, as the toxin encodes nuclease (CinB) and deubiquitinase (CidB) domains similar to RP toxins of certain wolbachiae that cause cytoplasmic incompatibility in arthropod hosts ^167–170^. We previously showed that pLbAR_38 shares similarity to a diverse assemblage of proteins from a narrow range of obligate intracellular bacteria, some of which are known reproductive parasites ^5^. Here, the remaining schema shows the result of a Blastp search against the NCBI nr database using pLbAR_38 (coordinates 1-3048, which excludes the ankyrin repeats) as the query (inset, color key for alignment scores). NOTE: each subject (numbered 1-24 at right) represents a distinct protein architecture, yet in some cases multiple similar proteins can be found for closely related species and strains). Yellow stars, novel RP toxins identified since our prior report ^5^. Matches for wolbachiae, *Cardinium* species (Bacteroidetes), *Diplorickettsia* species (*Gammaproteobacteria*), and *Rickettsiella* species (*Gammaproteobacteria*) are not shown to emphasize novel findings in Rickettsiaceae. White numbers indicate % identity across significant alignments. Subjects missing internal sequence relative to pLbAR_38 (no. 4) are joined by dashed lines; subjects with large insertions relative to pLbAR_38 are adjusted accordingly (nos. 5, 14, and 15). Blurred-out regions within subjects depict sequences with no significant matches to pLbAR_38. Six proteins for the *Rickettsia* endosymbiont of *Ceutorhynchus assimilis* are boxed, with the arrow pointing to a seventh protein that is directly compared to a protein from the *Rickettsia* endosymbiont of *Adalia bipunctata* (unpublished assembly), which contains the ovarian tumor (OTU) cysteine protease (Pfam OTU, PF02338) domain also found in the Spaid male-killer toxin of *Spiroplasma poulsonii* sp. ^171^. Inset, alignment of the OTU protease domains of select bacteria: TEOG, *Tisiphia* endosymbiont of *Oedothorax gibbosus*; RECA, *Rickettsia* endosymbiont of *Ceutorhynchus assimilis*; REAB, *Rickettsia* endosymbiont of *Adalia bipunctata*. Amino acid coloring is described in the **FIG. 3** legend.

**TABLE S1. RvhB4 sequences used for phylogeny estimation.**

**TABLE S2. Supporting information for REMs and cREMs.**

**TABLE S3. Supporting information for type VI secretion system analyses.**

**TABLE S3. Supporting information for patatin analyses.**

## Notes

### Competing Interest Statement

The authors have declared no competing interest.

